# Internal Model-based Modular Control Reproduces Multiple Lower-Limb Behaviours and Their Transitions Within a Single Simulation

**DOI:** 10.1101/2023.12.04.569887

**Authors:** David Muñoz, Donal Holland, Giacomo Severini

**Affiliations:** School of Electrical and Electronic Engineering, University College Dublin, Dublin, Ireland; School of Mechanical and Materials Engineering, University College Dublin, Dublin, Ireland

**Keywords:** Predictive modeling, musculoskeletal modeling, internal models, locomotion, muscle synergies

## Abstract

Computational models of neural control are a powerful tool for evaluating principles of motor control that cannot be tested directly in conventional experimental settings. Current gait controllers struggle with physiological grounding and can replicate a small set of discrete behaviours.

**Objective:** here we propose a new neural controller, the Internal Model-based Modular Controller, that can transition between multiple lower-limb motor tasks within a single simulation. The architecture comprises a simplified model of the Mesencephalic Locomotor Region, which activates different internal models. These internal models organise functional muscle synergies into task-specific networks that generate coordinated activity across multiple muscles.

**Results:** the controller performs Stand-To-Walk-To-Stand and Stand-To-Backward Walking-To-Stand transitions and modulates gait speed within a simulation by adjusting a few control signals. The observed simulated biomechanics and muscle activation patterns are generally consistent with experimental observations.

**Conclusions:** The proposed modular architecture is sufficient to produce heterogeneous motor behaviours and offers a testable hypothesis of their neural organisation.

**IMPACT STATEMENT:** This study presents a modular neural controller, which employs internal models and functional muscle synergies to reproduce heterogeneous legged behaviours within a single simulation framework.

## I. Introduction

PREDICTIVE neuromuscular models can address gaps in the knowledge of human motor control by testing hypotheses about the underlying neurophysiology [1]. In particular, models that include explicit formulations of neural control architectures can mimic the role of sensorimotor control in the generation of movements.

Several works in literature have proposed explicit formulations of the neural control of gait. Current controllers present architectures which normally include reflexes [2], [3], central pattern generators (CPG) [4], or a combination of both [5], [6], [7]. These architectures often present challenges related to dimensionality and physiological plausibility.

Reflex-based controllers implement reflex loops [2], [3], [8], [9], [10] to drive gait with remarkable similarity to experimental data. These controllers have shown that several principles of legged mechanics observed in gait (e.g., compliant leg behaviour, ankle push-off, hip unloading, swing, leg retraction) can emerge from relatively simple reflex architectures [2]. However, purely reflex-based controllers are often considered an oversimplified representation of neural control of gait as they restrict control to local sensorimotor feedback loops while neglecting supraspinal control, internal models, and context-dependent modulation that shape human locomotion [2], [3].

CPG-based controllers [4], [5], [6], [7], [11] incorporate these nominal structures as the primary element of control. CPGs are networks in the spinal cord that produce rhythmic locomotor activity even in the absence of higher-level control or sensory inputs [12], [13]. CPGs are typically implemented following a half-centre organisation [14]. This scheme describes a pair of antagonistic neurons (half-centres) coupled by mutual inhibitory disynaptic connections, forming a rhythmic unit. Gait control models based on the half-centre organisation can control the rhythm of motion using a few parameters, reducing control dimensionality [4], [15]. Both reflex and CPG architectures are better suited for simulations of isolated motor tasks and cannot fully explain the flexibility in lower limb control observed experimentally.

Modularity and hierarchy across components are features of physiological neural control that are often used to explain the experimentally observed flexibility in control [16]. Modules, or muscle synergies, are considered to be functional units of spinal interneurons that generate specific patterns of muscle activation [17], [18], [19]. The combination of a limited number of synergies in a hierarchical architecture results in different motor behaviours [17], [20] suggesting that modularity allows for few control signals to generate multiple motor outputs. Some neural controllers of gait [21] [5] have integrated synergies in a CPG-based controller. In these models the synergies were predetermined and periodically triggered by the CPG as bursts of activity, and were not used as modular blocks that can be combined to form different behaviors.

Here, we propose the Internal Model-based Modular Controller (IMMC) as a novel architecture that integrates functional synergies, mapping to the principles of legged mechanics, in a hierarchical system (Fig. 1) that can replicate multiple lower limb motions. The architecture aims to offer an explicit neural controller that can be used to simulate different lower limb behaviors in a physiologically plausible way. Beyond reproducing healthy locomotor behaviors, an explicit and interpretable representation of neural control is necessary for modelling pathological movement, which arises primarily from altered control rather than from altered musculoskeletal properties alone [22]. The IMMC could provide a basis for modelling control alterations.

**Figure 1.**
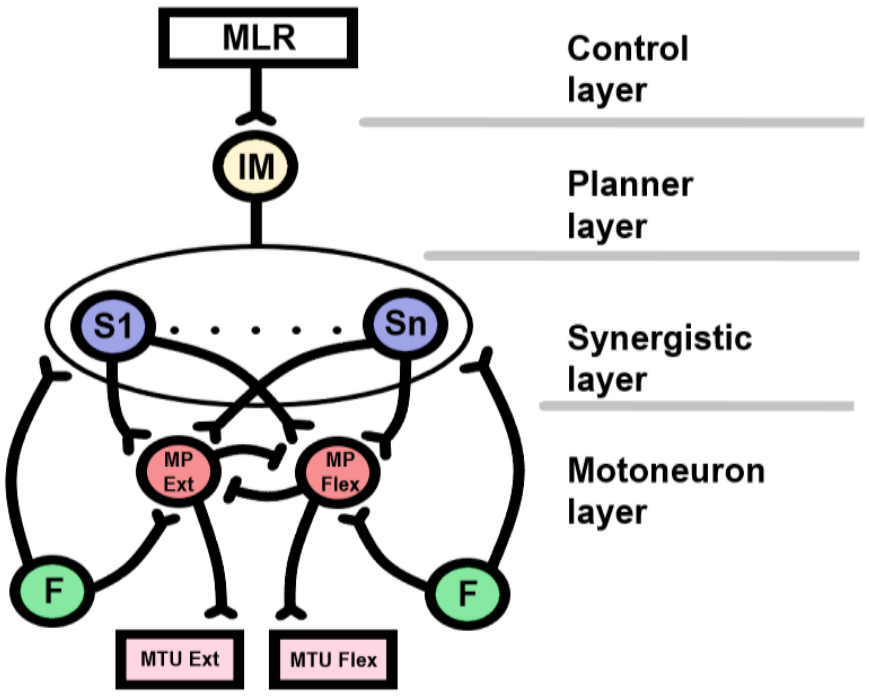
Scheme of IMMC, showing its hierarchy, from top to bottom: Control layer, Planner layer, Synergistic layer, and Motoneuron layer. MLR: mesencephalic locomotor region, IM: Internal Model, S1: first synergy, Sn: n-th synergy, MPExt/Flex: extensor/flexor motor pools, F: sensory feedback, MTUExt/Flex: extensor/flexor musculotendon unit.

The IMMC is based on five assumptions, inspired by previous literature (Table S4 in the Supplementary Materials summarises the evidence for each element and the modelling choices made):

i) motion can be mapped to an arrangement of functional synergies located in the spinal cord [20]; ii) some functional synergies have a reflexive nature [23]; iii) a task-specific network is the result of applying a specific pattern of excitatory and inhibitory synapses, here referred to as an internal model (IM), to a set of functional synergies [24]; iv) a hypothetical higher layer, corresponding to the mesencephalic locomotor region (MLR), activates IMs in parallel or sequentially according to the commanded task [25]; and v) standing, forward walking and backward walking tasks share synergies [26], [27], [28].

In the IMMC, an IM is the task-specific set of gains and between-synergies connections that turns an MLR command into a coordinated synergy network. It does not predict the sensory consequences of movement, as a forward model does, but maps a task onto the structure of the commands that achieve it, similarly conceptually to an inverse model [29].

The structure of the IMMC consists of four layers organised hierarchically (Fig. 1): 1) the Control layer, which activates the commanded task; 2) the Planner layer, which contains the IMs; 3) the Synergistic layer, which contains the synergies; 4) the Motoneuron layer which represents spinal reflexes and control signal integration.

Each synergy drives the motor pools through spatial gains (β) that set its amplitude and a temporal gain (θ) that sets its timing. The motor pools also receive sensory feedback through reflex gains (δ) and inhibit their antagonists through a gain (γ).

The output of this control architecture drives a simple biomechanical model to execute different tasks within a single simulation. We examine this capability across three transition scenarios: i) Stand-To-Walk-to-Stand (STWTS);
(ii) Stand-To-Backward Walk-To-Stand (STBWTS); (iii) forward gait speed transitions. Across these modalities, we assess the simulated motor dynamics (i.e., kinematics, muscle activations patterns, and ground-reaction forces) by comparing them with experimental data.

## II. Results

### A. Gait transitions

We first evaluated the capacity of the IMMC to reproduce the desired transition behaviours. In both the STWTS and STBWTS tasks the simulation begins with the CoM of the model in a static posture, and requires the model to start walking at t = 5 s, stop walking at t = 10 s and maintain a static posture for the remainder of the simulation (t = 15 s). For the speed transitions, the model again begins the simulation in a static posture, starts walking at t = 5 s and reaches the target speed by the end of the simulation (t = 10 s). Fig. 2 shows the results for all these transitions by showing the evolution of the antero-posterior velocity of the torso Centre-of-Mass (CoM) of the Head-Arms-Torso segment (HAT) across the full duration of the simulations.

**Figure 2.**
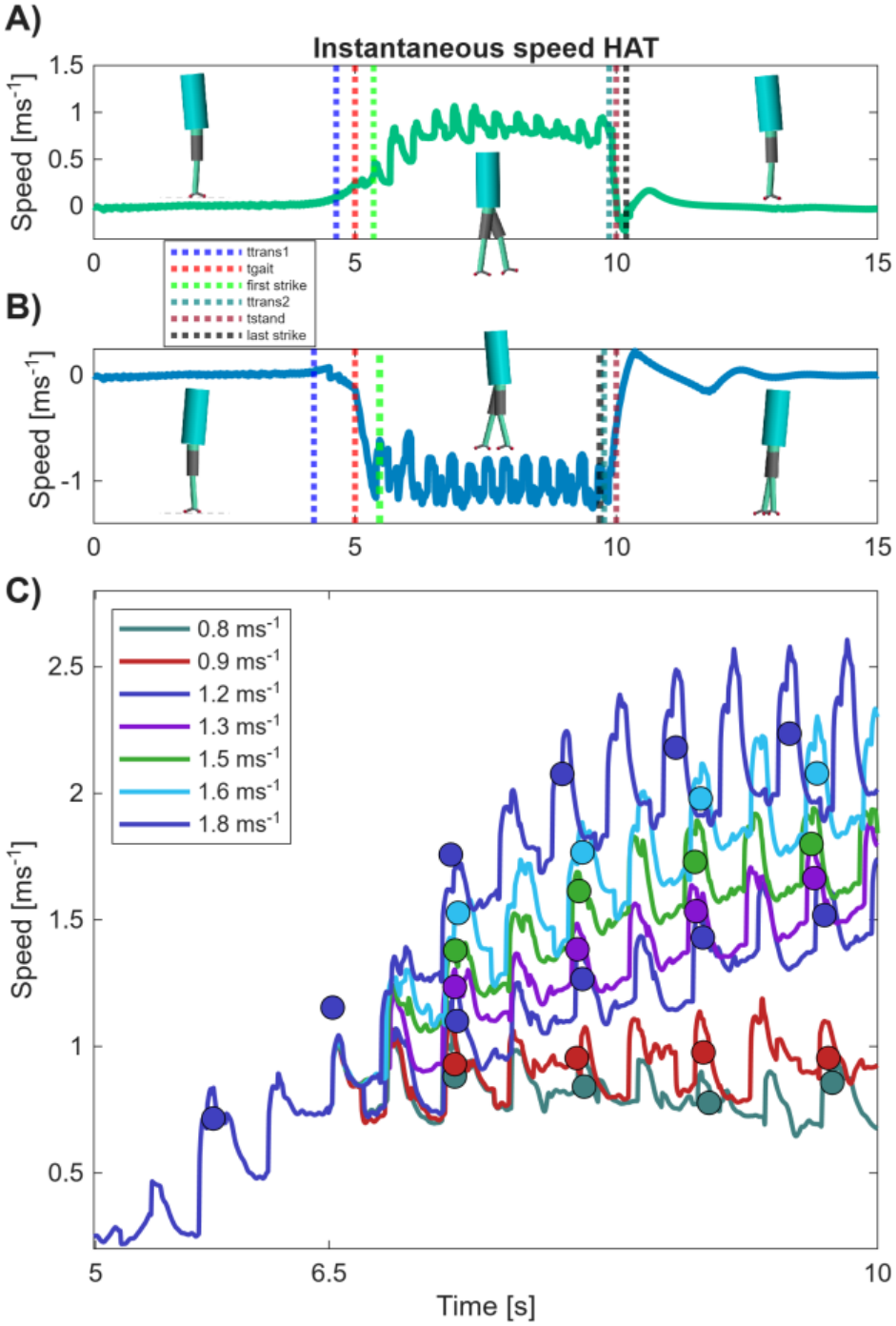
Instantaneous anterior/posterior speed of the Head-Arms-Torso (HAT). (A) Stand-To-Walk (t = 5 s) and Walk-To-Stand (t = 10 s) transitions at a gait speed of 0.8 ms^−1^. (B) Stand-To-Backward Walk (t = 5 s) and Backward Walk-To-Stand (t = 10 s) transitions. In A and B, markers show the onset of the transition internal model (t_trans1_, t_trans2_), the switches to gait (t_gait_) and to standing (t_stand_), the first heel strike after t_gait_ and the last foot contact before standing. (C) Speed transitions for forward gait at 0.8, 0.9, 1.2, 1.3, 1.5, 1.6 and 1.8 ms^−1^: instantaneous HAT speed (lines) and stride-averaged speed (circles). All speed simulations share the same gait initiation and diverge when the control signal is set to its target value at t = 6.5 s.

In the STWTS simulation (Fig. 2A) the controller maintains a stable standing in the first tract of the simulation. At the time of transition from stand to walk, t_trans1_, the model breaks the standing posture by leaning the trunk forward. Stepping starts at t_gait_ = 5 s and the oscillation of the CoM stabilises at t = 6.5 s. From that instant to t = 10 s, the model maintains a stable gait, with an average walking speed around 0.8 ms^−1^. At the time of transition from walk to stand, t_trans2_, after 11 steps, the CoM speed drops rapidly, and at t_stand_ = 10s, it begins to stabilise towards a quasi-static posture. This standing posture is successfully maintained until the end of the simulation (t = 15 s). The STBWTS simulation (Fig. 2B) presents the same standing dynamics as the forward condition until t_trans1_, when the model initiates a backward lean of torso. At t_gait_ = 5 s, backward stepping begins, and the CoM speed stabilises at −0.8 ms^−1^ by t = 6 s. A steady backward walking is then maintained until t_trans2_ when the model starts to decelerate, after 11 steps. At t_stand_ = 10 s the CoM speed stabilises toward zero, remaining in this quasi-static posture until t = 15 s. Fig. 2A and B mark the first heel strike after *t*_gait_ and the last foot contact before standing in both simulations.

The IMMC can achieve speed transitions by modulating the control signal for forward gait. Fig. 2C shows the instantaneous speed of CoM for the seven tested speed transitions (0.8, 0.9, 1.2, 1.3, 1.5, 1.6, and 1.8 ms^−1^). The speed is set by the control signal of the forward-walking IM, *u*, which modulates its gains (*u* = 1 at 1.3 ms^−1^; Supplementary Eqs. 1–2) and ranged from *u* = 1.6 at 0.8 ms^−1^ to *u* = 0.4 at 1.8 ms^−1^ (Table S5). The simulations share the same gait initiation and diverge when *u* is set to its target value at *t* = 6.5 s, so the change of speed occurs during walking. Over the walking period, the mean stride speed was within 8 % of the target speed, but at 1.2 ms^−1^ and above it was still increasing at *t* = 10 s (Fig. 2C; Table S6). In longer simulations (up to 20 s of walking, Fig. S5), these speeds settled 0.3–0.5 ms^−1^ above their targets (Table S6). The metabolic cost of transport for each speed is shown in Fig. 3. For the speed of 1.3 ms^−1^, close to the preferred walking speed by humans [30], [31], the average cost of transport is also close to that observed experimentally (3.5 Jkg^−1^m^−1^) [32], [33]. The cost of transport was highest at the slowest speeds (Fig. 3 and Supplementary Material Results A).

**Figure 3.**
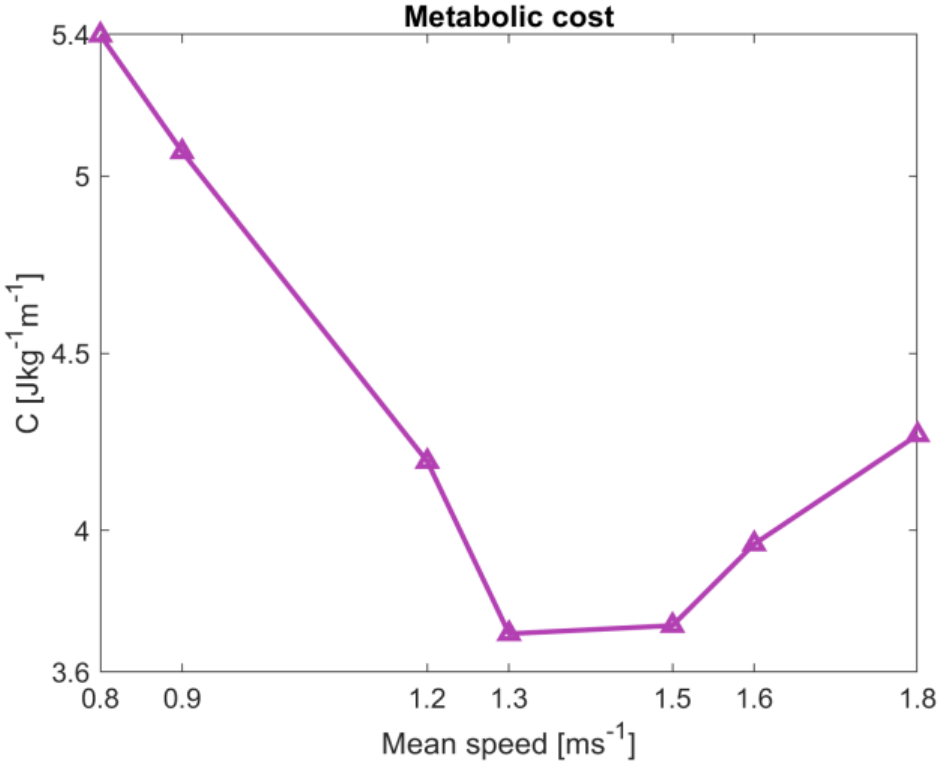
Metabolic cost of all the forward gait speeds.

### B. Gait biomechanics

Figure 4 compares the simulated joint trajectories and muscular activations of forward walking with experimental data [28]. The simulation used for this analysis was the speed transition with target speed of 1.3 ms^−1^, and the data represent the average calculated across 10 gait cycles (five per leg) of steady walking, each segmented between consecutive heel strikes (Supplementary Material Section I). Results were evaluated by calculating cross-correlation coefficients (*r*) and the corresponding lag (*Δ*) between the simulated and experimental traces, together with the root-mean-square error (RMSE) and the differences in peak magnitude and peak timing (Table 1).

**Table 1.**
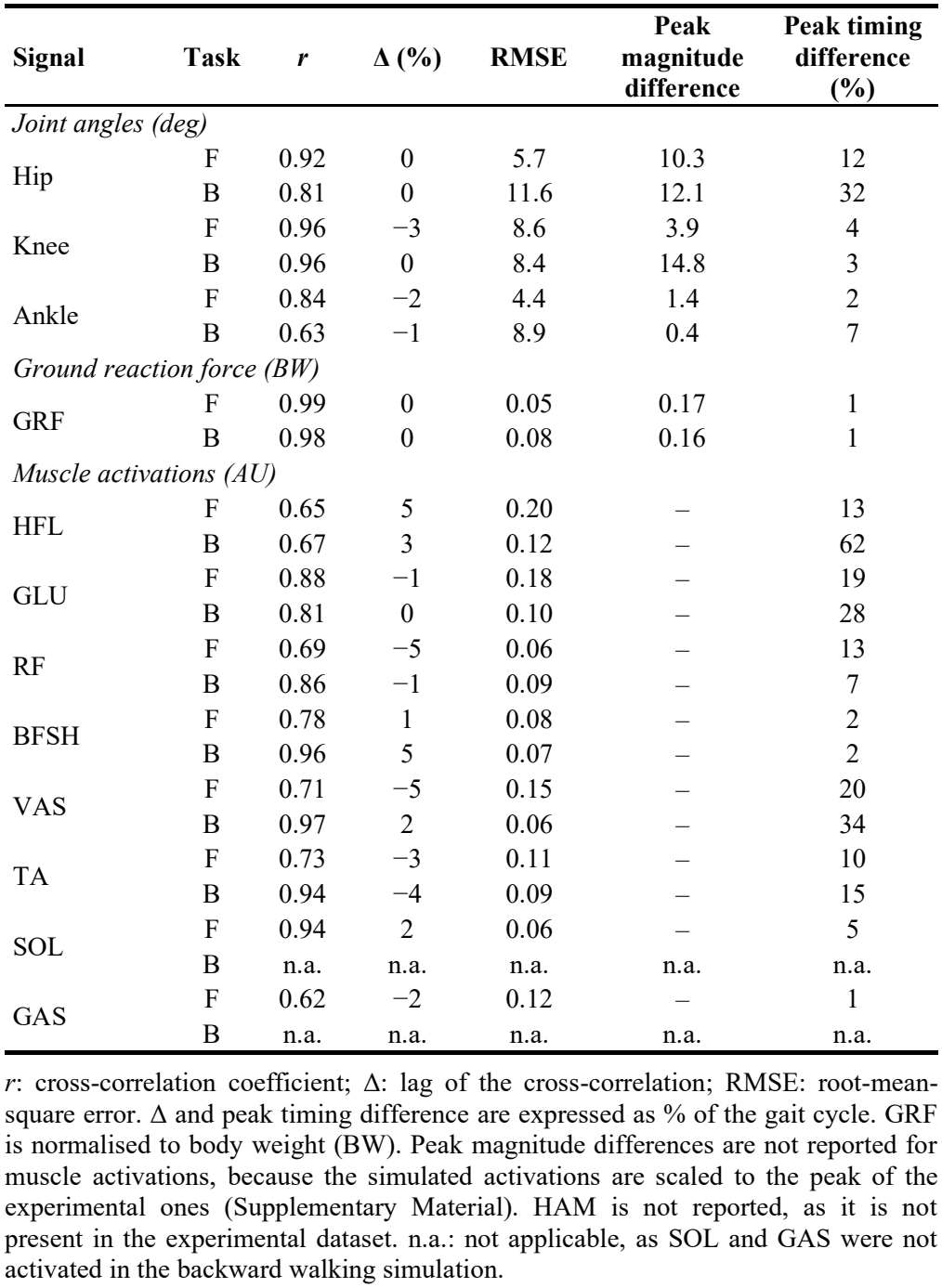
Comparison between simulated and experimental data for forward walking (F, 1.3 ms^−1^, Fig. 4) and backward walking (B, 0.8 ms^−1^, Fig. 5). Experimental data from [28].

| **Signal** | **Task** | ***r*** | **Δ (%)** | **RMSE** | **Peak magnitude difference** | **Peak timing difference (%)** |
|---|---|---|---|---|---|---|
| *Joint angles (deg)* |  |  |  |  |  |  |
| Hip | F | 0.92 | 0 | 5.7 | 10.3 | 12 |
|  | B | 0.81 | 0 | 11.6 | 12.1 | 32 |
| Knee | F | 0.96 | −3 | 8.6 | 3.9 | 4 |
|  | B | 0.96 | 0 | 8.4 | 14.8 | 3 |
| Ankle | F | 0.84 | −2 | 4.4 | 1.4 | 2 |
|  | B | 0.63 | −1 | 8.9 | 0.4 | 7 |
| *Ground reaction force (BW)* |  |  |  |  |  |  |
| GRF | F | 0.99 | 0 | 0.05 | 0.17 | 1 |
|  | B | 0.98 | 0 | 0.08 | 0.16 | 1 |
| *Muscle activations (AU)* |  |  |  |  |  |  |
| HFL | F | 0.65 | 5 | 0.20 | – | 13 |
|  | B | 0.67 | 3 | 0.12 | – | 62 |
| GLU | F | 0.88 | −1 | 0.18 | – | 19 |
|  | B | 0.81 | 0 | 0.10 | – | 28 |
| RF | F | 0.69 | −5 | 0.06 | – | 13 |
|  | B | 0.86 | −1 | 0.09 | – | 7 |
| BFSH | F | 0.78 | 1 | 0.08 | – | 2 |
|  | B | 0.96 | 5 | 0.07 | – | 2 |
| VAS | F | 0.71 | −5 | 0.15 | – | 20 |
|  | B | 0.97 | 2 | 0.06 | – | 34 |
| TA | F | 0.73 | −3 | 0.11 | – | 10 |
|  | B | 0.94 | −4 | 0.09 | – | 15 |
| SOL | F | 0.94 | 2 | 0.06 | – | 5 |
|  | B | n.a. | n.a. | n.a. | n.a. | n.a. |
| GAS | F | 0.62 | −2 | 0.12 | – | 1 |
|  | B | n.a. | n.a. | n.a. | n.a. | n.a. |

*r*: cross-correlation coefficient; Δ: lag of the cross-correlation; RMSE: root-mean-square error. Δ and peak timing difference are expressed as % of the gait cycle. GRF is normalised to body weight (BW). Peak magnitude differences are not reported for muscle activations, because the simulated activations are scaled to the peak of the experimental ones (Supplementary Material). HAM is not reported, as it is not present in the experimental dataset. n.a.: not applicable, as SOL and GAS were not activated in the backward walking simulation.

**Figure 4.**
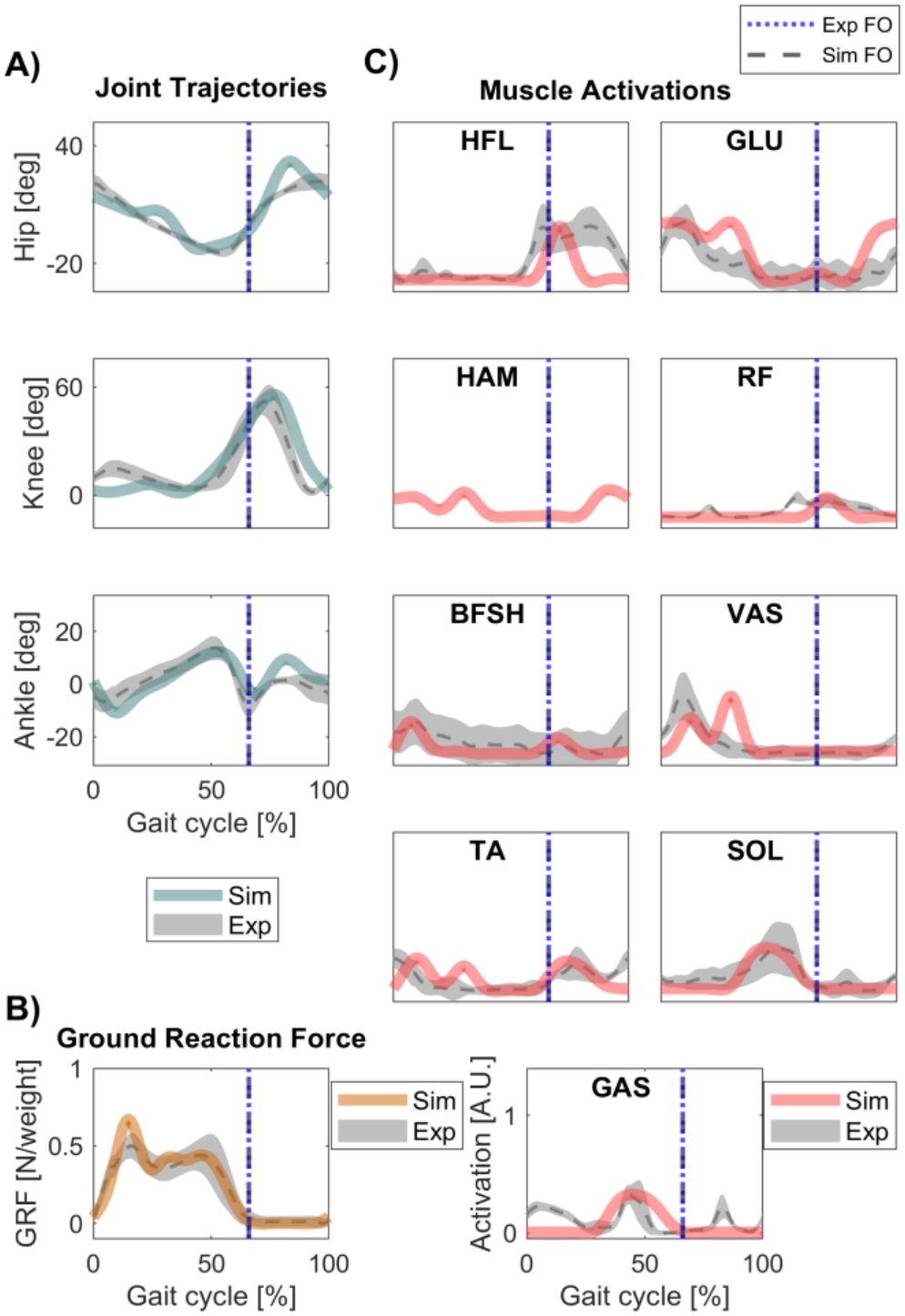
Comparison between simulated and experimental biomechanics and muscular activation for forward walking. (A) Experimental and simulated angular positions for hips, knees, and ankles at 1.3 ms^−1^. Green: simulated data, grey: experimental data. Upward direction: flexion, downward direction: extension. (B) GRF across the gait cycle at 1.3 ms^−1^. Experimental and simulated GRF. Yellow: simulated data, grey: experimental data. (C) Muscle activation patterns along the gait cycle at 1.3 ms^−1^. Orange: simulated data, grey: experimental data. Experimental data for HAM was not present in the referenced dataset. Experimental angular positions, GRF, and muscle trajectories taken from [28]. Blue vertical dotted line: experimental foot-off (66 %), black vertical dotted line: simulation foot-off (66 %).

Foot-off occurs at 66 % of the gait cycle in both the simulation and the experimental data. The simulated hip, knee, and ankle angle are consistent with the experimental trajectories (Fig. 4A), with high cross-correlation coefficients (*r* = 0.84–0.96; Table 1). Despite the strong correlations, hip exhibits more extension than its experimental counterpart at the beginning and end of the gait cycle (peak difference of 10.3°). The simulated ankle is also in a slightly more dorsiflexed position at the beginning and end of the gait cycle. The GRFs (Fig 4B) match the typical pattern of contact force during walking [28], with a high cross-correlation (*r* = 0.99). The peak associated with weight acceptance in the simulated GRF is slightly larger than its experimental counterpart.

Fig. 4C presents the comparison between simulated and experimental muscle activation patterns across the gait cycle for the hip flexors (HFL), gluteus (GLU), hamstrings (HAM), rectus femoris (RF), biceps femoris short head (BFSH), vastus (VAS), tibialis anterior (TA), soleus (SOL), and gastrocnemius (GAS). The hamstrings (HAM) muscle was present in the simulations but not in the experimental dataset. Overall, the simulated activation timing falls within the experimentally observed ranges reported by Zych et al. [28]. Cross-correlation coefficients ranged from 0.62 for GAS to 0.94 for SOL (Table 1). The lowest similarity was found for GAS, whose pattern poorly matched its experimental counterpart, RF, which was active only during the first half of the gait cycle, and HFL, whose activity was generally reduced along the cycle.

Fig. 5 shows the comparison of the simulated trajectory of the joints over the backward gait cycle to the angular positions in humans [28]. The simulated data used in this analysis were the ones relative to the walking portion of the STBWTS simulation. The speed of backward walking was selected to be 0.8 ms^−1^, within the range of preferred backward velocities [34].

**Figure 5.**
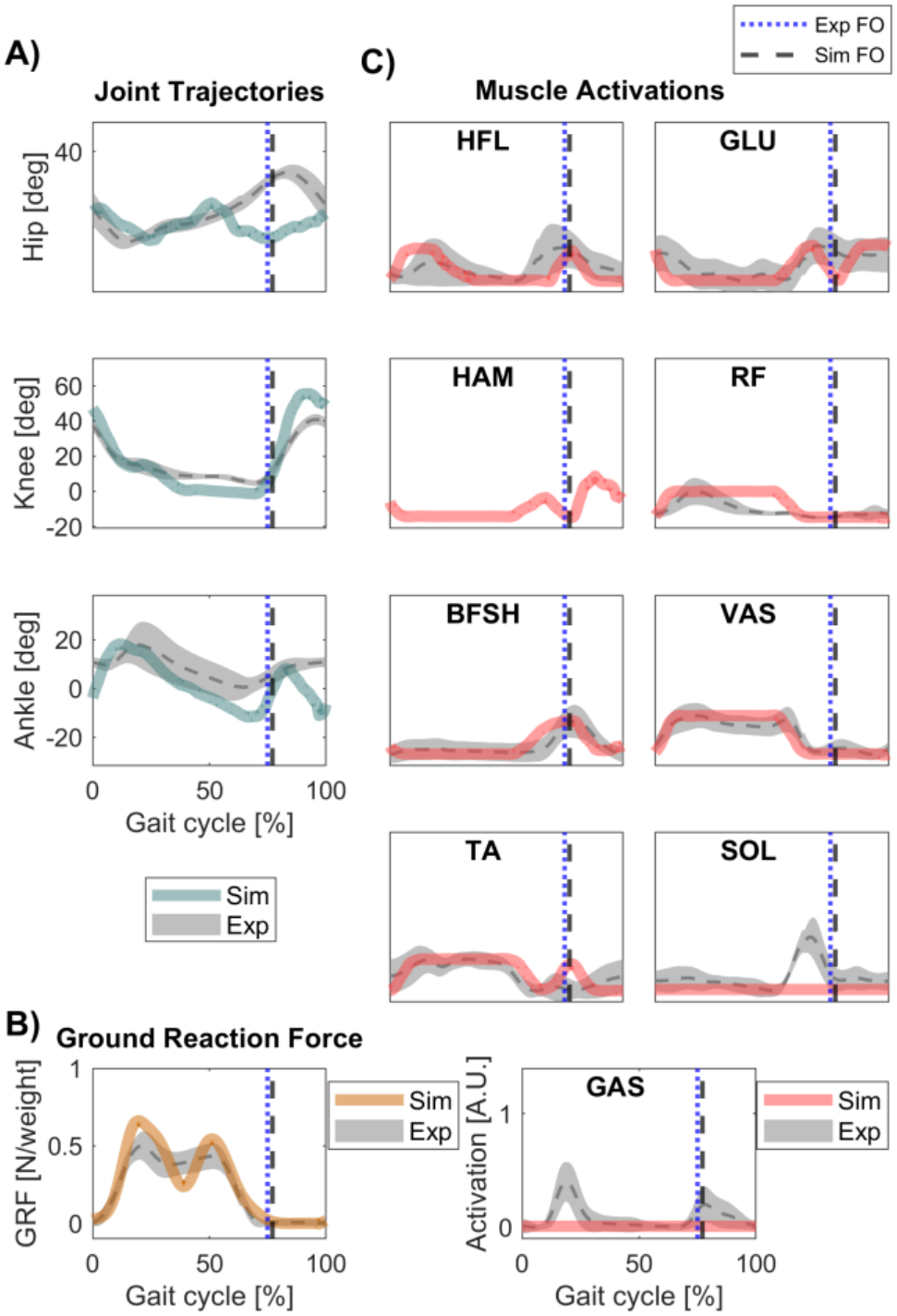
Comparison between simulated and experimental biomechanics and muscular activation for backward walking. (A) Experimental and simulated angular positions for hips, knees, and ankles at 0.8 ms^−1^. Green: simulated data, grey: experimental data. Upward direction: flexion, downward direction: extension. (B) GRF across the gait cycle. Experimental and simulated GRF. Yellow: simulated data, grey: experimental data. (C) Muscle activation patterns along the gait cycle. Orange: simulated data, grey: experimental data. Experimental data for HAM was not present in the referenced dataset. Experimental angular positions, GRF, and muscle trajectories taken from [28]. Blue vertical dotted line: experimental foot-off (76 %), black vertical dotted line: simulation foot-off (77 %).

The simulation foot-off occurs at 77 % of the gait cycle, while the experimental foot-off occurs at 76 %. The simulated hip, knee, and ankle angles are consistent with the experimental trajectories (*r* = 0.63–0.96; Table 1). The simulations presented increased hip extension in the second half of the cycle (peak difference of 12.1°), and increased ankle plantarflexion at the beginning and at the end of the cycle. The largest peak difference was at the knee (14.8°; Table 1). Fig. 5B shows the GRFs. The results match the typical pattern of contact force during walking [28], with a good cross-correlation (*r* = 0.98). The simulated muscular activation timing (Fig. 5C) falls within the experimentally observed ranges reported by Zych et al [28], with the clear exception of SOL and GAS which are not excited in the backward gait simulations. For the remaining muscles, cross-correlation coefficients ranged from 0.67 for HFL to 0.97 for VAS (Table 1). The joint with lowest similarity is the ankle, which is excessively plantarflexed. HFL also presents the highest disagreement due to the activity decrement in the second half of the cycle. Muscle synergies extracted from the simulated activations matched the experimental synergies of forward walking (cosine similarity 0.90–0.94) and one of the synergies of backward walking. The other synergies include SOL and GAS and were matched only in part (Supplementary Material, Fig. S6 and Table S8). Comparison of the biomechanics and muscular activations for the speed transitions is presented in the Supplementary materials (Fig. S4).

## III. Discussion

The IMMC achieved all target behaviours and can achieve transitions between standing and both forward and backward walking within the same simulations. We also demonstrate that the controller can walk at different speeds. The operational workflow of the proposed hierarchical architecture is simple: the Control layer activates the internal models of the commanded tasks, each mapping to a shared set of functional synergies. Each internal model organises the synergies within a task-specific network, with a different flow of synergies control between forward and backward walking (Fig. S2). Backward walking uses synergies gains up to an order of magnitude larger than forward walking. Standing and forward walking share the gains of S1 and differ in its timing (θ). The gains of each IM are listed in Table S1.

The controller requires only a small number of control signals to drive transitions between tasks. As shown above, forward speed modulation can be achieved through a single control signal. The controller’s ability to reconfigure a limited number of motor primitives to generate distinct tasks is consistent with experimental evidence on modular motor control [26], [27], [28], [35].

The simulation results generally agree with the experimental biomechanics reported for forward and backward walking [28] and for gait at different speeds [36] (see Supplementary materials), and most of the discrepancies we observed can be partially attributed to the simplicity of the biomechanical model employed for the simulations, which also limits the synergistic and reflex structure (e.g torso-related muscles and vestibular reflexes are missing in our implementation [37]). Abnormal ankle trajectories are common in predictive simulations driven by neural control [3], [5], [7], [39] and Schumacher et al. [38] showed that abnormal ankle trajectories often depend on the complexity of the musculoskeletal model.

Muscle activation timing also agreed with the experimental data for most muscles. The main exceptions were HFL, RF and GAS in forward walking, and SOL and GAS in backward walking, where the two plantarflexors were not activated. The GAS discrepancies likely reflect its implementation as a scaled copy of SOL activity, which cannot reproduce the parts of its experimental pattern that differ from SOL, while RF, which sums the hip flexor and knee extensor pools, is dominated by the knee extensor activity of early stance. In backward walking, the cost function (Supplementary Eq. 15) penalises metabolic cost but not joint-limit torques, so the optimiser could keep the ankle passively at its limit instead of activating the plantarflexors. However, the simulation still reproduced the experimental joint angles with fidelity. The planar model can therefore match joint kinematics with different muscle strategies.

The architecture of the IMMC was defined a priori, and only its gains were optimised. Our results show that it is sufficient to generate standing, forward and backward walking, speed modulation and the transitions between them. They do not establish that the nervous system uses the same organisation, and other arrangements of modules or triggers may also obtain similar results. The extended and time-shifted simulations also show that the behaviours are robust around the durations and command timings used in the optimisation (Supplementary Results B). We did not assess how sensitive the results are to the controller parameters or to the architectural assumptions summarised in Table S4. Because each assumption is a separate element of the controller, future work can modify and test each one independently, guided by the muscle-level mismatches observed here, for example through different motor-pool mappings or cost functions (e.g. a joint-limit penalty or a term penalising fully inactive muscles [7]).

In these simulations, the IMMC executes transitions but does not select which transition occurs or when. For simplicity, the control signals follow a fixed schedule: the walking IM is activated at *t* = 5 s and the standing IM at *t* = 10 s, and the optimisation found the onset of the transition IM before each switch (*t*_trans_). Within this schedule, the transition synergy, modulated by trunk lean, breaks quiet standing before the first step, or slows the body and brings the feet together before standing. The time-shifted simulations show that the transitions are completed mainly within this schedule (Table S7). Thus the current implementation of the IMMC does not model how the MLR selects and times transitions, for example from goals or sensory events.

A common limitation of neural controllers of gait is their task specificity, so that most explicit formulations of neural control of gait have either been tested on a single task or, when able to replicate different tasks, they require ad-hoc optimizations for each of them. We recently showed that a hybrid CPG-reflexes architecture can reproduce both walking and cycling, although through distinct optimisation of the controller parameters [7]. Similarly, Bunz and colleagues showed that a simple reflex controller can replicate different lower limb tasks including forward and backward walking, running and hopping [40]. Also in that case, the controller required different optimisation of the control parameters for each task. Works in literature have shown control architectures capable of replicating different tasks in the same simulations, by sequentially engaging distinct controllers. For example, Van der Kruk & Geijtenbeek [10] developed a reflex-based Sit-To-Walk model that integrated distinct sit-to-stand and gait controllers. Ramadan and colleagues [39] used internal models in a reflex controller to switch between forward gait patterns within a simulation, keeping the lower-level parameters fixed and selecting or interpolating higher-level parameters from a library of pre-optimised gaits. With respect to this work, the IMMC shares the principle of a fixed lower layer modulated from above, but its higher layer changes the structure of the network rather than only its parameters. Each IM reorganises the same functional synergies into a different network, changing their order and coupling as well as their gains. This allows the same synergies and reflex pathways to produce qualitatively different tasks (standing, forward walking and backward walking), together with dedicated transitions that start and stop gait from quiet standing. Once optimised, the IMMC is controlled only through the signals that activate the IMs and set the forward speed.

The explicit and modular structure of the IMMC may also be relevant to the study of movement impairments. Personalised neuromusculoskeletal models have been proposed as a means of predicting and optimising the functional outcome of clinical interventions in stroke, osteoarthritis, and other neurological conditions [22]. A recurring conclusion of this line of work is that the neural control of movement, and not only its musculoskeletal substrate, must be represented explicitly, since many impairments are best described as alterations of the feedforward drive or of healthy versus unhealthy feedback. In these frameworks, muscle synergies are typically extracted from EMG through non-negative matrix factorisation and then employed as control primitives. However, such synergies are emergent, descriptive quantities that merge the feedforward and feedback processes that generate them [41]. In the IMMC the functional synergies are primitives organised by internal models, with feedforward commands and feedback modulation represented as separate pathways. This separation could allow specific impairments to be mapped onto distinct control elements, potentially bridging the synergies recovered through factorisation and the mechanisms that produce them. Realising this potential would require personalisation to individual data and a more detailed biomechanical model, but the architecture provides a principled starting point for such extensions.

Overall, the IMMC is a hierarchical controller in which standing, forward walking and backward walking are produced by task-specific IMs acting on a shared set of functional synergies, without switching between separate controllers. It provides a computational implementation of the hypothesis that locomotor versatility arises from the reuse and reweighting of shared control components across tasks, and a framework in which that hypothesis can be tested further.

## IV. Materials and Methods

### A. Controller Architecture

The structure of the IMMC consists of four layers organised hierarchically. From higher to lower, the layers are: 1) Control layer, representing a simplified model of the MLR; 2) Planner layer, constituted by the internal models (IMs), each one representing a specific motor task; 3) Synergistic layer, comprising six distinct functional synergies (S), each contributing to a specific phase of locomotion; 4) Motoneuron layer, where motor pools integrate the inputs from the functional synergies and sensory afferents before activating the MTUs. The propagation of the control signals within and across layers is regulated by sets of gains (β, θ, γ, and δ) that are found during the optimisation process.

The Synergistic layer comprises six distinct functional synergies: S1) *Compliant-leg-behaviour* resists gravity by extending the lower limbs; S2) *Ankle push-off* generates forward propulsion of the body; S3) *Hip unloading* reduces weight support on the stance leg and facilitates swing; S4) *Swing* propels the swing leg forward; S5) *Leg retraction* decelerates the swing and positions the leg for landing; S6) *Transition*, an ad hoc functional synergy that eases the transition between motor tasks. The first five synergies correspond with the principles of legged mechanics [2] and each contributes to a specific phase of locomotion (Figure 6), while the sixth synergy is used to control transitions between gait (forward and backward) and stand and vice versa. The control flow of the IMMC and the equations and control rules driving each layer are fully explained in the Supplementary material (Fig. S1, S2 and S3).

**Figure 6.**
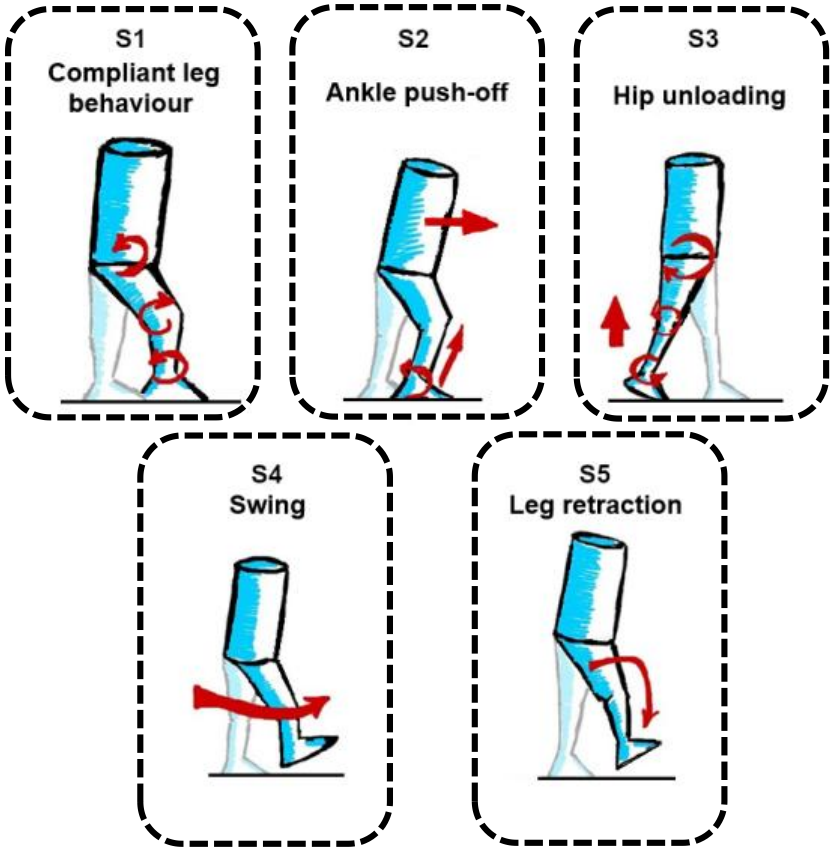
Principles of the legged mechanics. The figure presents the five principles of legged mechanics. The synergies implemented in the IMMC are based on these principles.

### B. Optimisation procedure

The objective of the optimisation is to tune the parameters characterising the IMMC. The optimisation was carried out in six stages, each corresponding to a task: 1) standing task; 2) transition to forward walking at 1.3 ms^−1^ (STW); 3) transition to forward walking at 0.8, 0.9, 1.2, 1.5, 1.6, and ms^−1^; 4) transition from 0.8 ms^−1^ walking to standing (WTS); 5) transition to backward walking (STBW); 6) transition from backward walking to standing (BWTS). A total of 163 parameters were optimised and 15 were hand-tuned. The code of the IMMC and the final values and optimisation bounds of all parameters are available at https://github.com/davidmunozrodr/IMMC. Each stage optimises the parameters of the corresponding internal models. Stage 1 and 2 also optimises the parameters of the system that are general to the architecture and do not belong to any internal model (See Supplementary Materials). To ease the optimisation process, the initial guesses for the parameters driving forward and backward gait (stage 2 and 5) were hand-tuned. Once the parameters were optimised, they were kept fixed within the IMMC. The algorithm used in all optimisations is the Covariance Matrix Adaptation Evolution Strategy (CMA-ES) [42]. Each stage is optimised with a maximum of 600 iterations, 5 parents, and 40 offspring in each generation. The optimization procedure used ad-hoc cost functions for each behavior (and thus IM), described in the Supplementary materials.

## Supplementary Materials

The Supplementary material presents extended details on the biomechanical model used in the analysis, the architecture of the controller and the optimisation procedure. We also present the additional details on the results, including the biomechanical and muscular activations comparisons for the speed transitions.

