## Supplementary Materials, Figure S1, Table S1, Figure S2, Table S2, Figure S3, Table S3, Table S4, Figure S4, Table S5, Table S6, Figure S5, Table S7 for "Internal Model-based Modular Control Reproduces Multiple Lower-Limb Behaviours and Their Transitions Within a Single Simulation"

David Muñoz, Donal Holland, and Giacomo Severini*., *Senior Member, IEEE*

THE Supplementary Materials presents an in-depth description of the Methodology of the IMMC controller with supplementary results relative to the speed transitions simulations.

#### I. Supplementary Methods

##### *A. Musculoskeletal model*

The musculoskeletal model was implemented following the proportions detailed in Song and Geyer [1] but constraining motion in the sagittal plane. This configuration was chosen for its simplicity and suitability for testing an initial implementation of the IMMC. The model represents a human body of 1.80 m height and 80 kg weight. It has seven segments representing HAT (head-arms-torso), thighs, shanks, and feet. The model has 6 degrees of freedom (DoF) as revolute joints connecting the segments, hips, knees, and ankles. Three additional DoFs were added to define motion with respect to the ground (longitudinal translation, vertical translation, and pitch rotation). The muscle layer is composed of nine Hill-type musculotendon units (MTUs) per leg: gluteus (GLU), hip flexors (HFL), hamstrings (HAM), rectus femoris (RF), vastus (VAS), biceps femoris short head (BFSH), gastrocnemius (GAS), soleus (SOL), and tibialis anterior (TA). Contact geometries consist of two spheres of 4 cm radius placed at the toes and heels of each foot (four in total). Sensory delays were added according to the position of the sensory source in the musculoskeletal model. Sources placed in a more distal position present a longer delay as they are further from the spinal cord. They are short, medium, or long (5, 10, 20 ms), according to their proximity to HAT/hips, knees, and ankles, respectively. They are denoted by $\Delta t_5$, $\Delta t_{10}$, and $\Delta t_{20}$. This implementation follows Geyer's models [2], [3]. These delays apply for all the sensory sources in the controller, including those related to trigger and modulate the functional synergies. The biomechanical model, the IMMC, optimisation and subsequent data analysis were implemented on Simulink/MATLAB R2022a.

##### *B. IMMC Control flow*

The MLR activates a specific internal model (Fig. S1), enabling the corresponding motor task, through control signals, $u$. The internal model works de-facto as a sub-circuit that organises and regulates the activity of the different synergies.

Each synergy projects to all motor pools, through specific gains. The motor pools integrate the activation from the different synergies and from joint-specific reflex pathways to derive the final activation to the muscles.

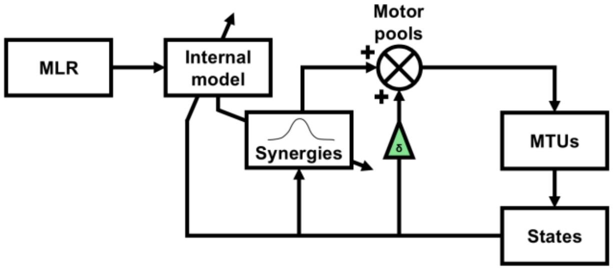


Figure S1. General workflow of the IMMC. Colours: $\delta$ gains (green)

The interaction between the MLR, the internal models, and the functional synergies can be interpreted as an open-loop control pathway. Together, these three components generate feedforward motor commands to achieve a specific objective within the broader motor task. For example, the compliant-leg-behaviour synergy (S1) prevents leg collapse during the stance phase of forward walking. However, these three blocks are not purely open-loop, because the functional synergies can be both triggered and temporally modulated by sensory signals through the internal models (see The Planner Layer).

##### *C. The Control Layer*

The Control layer is the highest hierarchical structure of the IMMC. The role of the Control layer in IMMC is that of controlling movement initiation and triggering the different internal models [4], [5]. The Control layer sends a limited number of control signals, denoted by $u$, that selects the active internal model. Generally, only one internal model is active during a task, with the exclusion of the transition internal model that is triggered concurrently to another internal model to facilitate the transition to the next task. Forward gait, backward gait, and standing are activated by a specific control signal, namely $u_{gait}$, $u_{back}$, and $u_{stand}$, respectively. While $u_{back}$, and $u_{stand}$ can take 0 (inactive) or 1 (active) values, $u_{gait}$ can take multiple non-zero values to modulate speed. The transition internal model can be activated by four control signals, one for each transition behaviour: $u_{STW}$ (Stand-To-Walk), $u_{WTS}$ (Walk-To-Stand), $u_{STBW}$ (Stand-To-Backward Walking), and $u_{BWTS}$ (Backward Walking-To-Stand). Each of these control signals can only take 0 (inactive) and 1 (active) values. In the simulations presented here, the timing of the control signals is set externally (Section G), so the Control layer does not decide which transition occurs or when.

### D. *The Planner Layer*

The Planner layer stores the internal models. The internal models have three roles: 1) regulate the temporal activation of the synergies (through the $\theta$ gains); 2) modulate the amplitude of the synergies (through the $\beta$ gains); 3) Interconnect the synergies in networks, where each synergy influences the activation of the others. In total, the IMMC presents four internal models, one for each task: forward gait, backward gait, standing and transition.

Forward and backward gait utilise functional synergies related to different principles of legged mechanics (see Synergistic layer). In both tasks, each synergy has six $\beta$ gains, linking each synergy to each motor pool (mapping to the flexion/extension of the three joints), and one $\theta$ gain. The set of $\beta$ and $\theta$ gains is specific to each internal model, except for the β gains of S1, which are shared by the standing and forward-walking internal models. The internal model for the standing task utilises a single functional synergy, the *compliant-leg-behaviour synergy* (S1, see the Synergistic Layer) and the corresponding internal model just provides six $\beta$ gains and one $\theta$ gain. The transition internal model also recruits only one functional synergy, *transition synergy* (S6), governing the four possible transitions: 1) from standing to forward walking (STW), 2), from forward walking to standing (WTS), 3) from standing to backward gait (STBW), and 4), from backward gait to standing (BWTS). As a result, the internal model contains one set of six $\beta$ gains and one $\theta$ gain per transition. The β and θ gains of each internal model are listed in Table S1.

Table S1. Synergy order, β gains and θ gains of each internal model.

| Synergy | β gain to motor pool *j* | | | | | | θ |
|---|---|---|---|---|---|---|---|
| | 1 | 2 | 3 | 4 | 5 | 6 | |
| *Standing* | | | | | | | |
| S1 | 1.3628 | −0.0003 | 2.5378 | 1.5453 | −1.5147 | 0.4722 | 0 / 0.25 |
| *Forward (u = 1)* | | | | | | | |
| S1 | 1.3628 | −0.0003 | 2.5378 | 1.5453 | −1.5147 | 0.4722 | −0.7750 |
| S2 | 1.7985 | −2.3853 | −0.8972 | −5.2753 | 1.0431 | 0.6737 | 0.4737 |
| S3 | −0.4297 | 1.9274 | −0.1852 | −0.3461 | 1.2412 | 1.8528 | −0.8061 |
| S4 | −0.1371 | 1.1577 | 0.5346 | 1.0112 | −1.1507 | 2.2536 | −0.4196 |
| S5 | 1.3762 | −0.6232 | 1.4603 | 0.7456 | −0.9802 | 0.8317 | −0.3750 |
| *Backward* | | | | | | | |
| S1 | −1.4657 | 12.3392 | 22.7519 | −6.1146 | 12.5565 | 13.7406 | −2.7318 |
| S5 | 0.3884 | 0.3314 | 0.0553 | −1.0300 | −0.6203 | 0.0692 | 9.3365 |
| S3 | −6.0246 | 12.7376 | −18.9190 | 27.9194 | −4.6450 | −0.5369 | 4.6624 |
| S4 | 2.1399 | −2.0832 | −1.6528 | −1.4289 | −1.5678 | −2.9801 | 6.7849 |
| S2 | −6.0494 | −6.9916 | 14.3385 | −0.7386 | −1.0948 | 3.5908 | −1.8807 |
| *Transition* | | | | | | | |
| S6 (STW) | 0.8516 | 1.5474 | −2.5641 | −4.5047 | −1.7062 | −0.4886 | 0 / 0.0433 |
| S6 (WTS) | −5.0068 | 4.5579 | 0.6195 | −4.7456 | −2.0446 | 21.3919 | 0 / 0.25 |
| S6 (STBW) | −2.2448 | −5.0111 | −0.5101 | −8.5323 | −0.0192 | −3.4698 | 0 / 0.25 |
| S6 (BWTS) | −0.5891 | 0.1522 | −0.6307 | −0.0397 | 0.3566 | −0.4053 | 0 / 0.25 |

Synergies are listed in the order of the network of each internal model; the backward network swaps S2 and S5. β: gain from the synergy to motor pool *j* (*j* = 1–6: hip extensor, hip flexor, knee extensor, knee flexor, ankle extensor, ankle flexor). For forward walking, the gains correspond to $u = 1$ (1.3 $ms^{-1}$); the gains at the other speeds follow from Eqs. 1–2 and the values of $u$ in Table S5.

The β gains of S1 are shared by the standing and forward-walking internal models. The values and optimisation bounds of all parameters are listed in the file IMMC_parameters.csv, available with the code at https://github.com/davidmunozrodr/IMMC.

The activation of the different internal models depends on the control layer. This layer sends a control signal, denoted by $u$, to the internal models. If $u = 0$, the internal model will remain inactive, and the synergies will not undergo a modulation process. If $u = 1$, the internal model will be activated, applying specific modulations to the synergies. In the case of forward walking, where different gait speeds were modelled, the activated state is not assigned to $u = 1$. Instead, $u > 0$ indicates an activated state, and different values of $u$ modulates the speed of gait. This is achieved by setting the β and $\theta$ gains as polynomial functions of $u$:

$$\beta_{ij} = \hat{\beta}_{ij} + \sum_{p=1}^{n} \hat{\beta}_{ip}(1-u)^p \tag{1}$$

$$\theta_i = \hat{\theta}_i + \sum_{p=1}^{n} \hat{\theta}_{ip}(1-u)^p \tag{2}$$

where $\beta_{ij}$ is the gain relating the $i$-th synergy with the $j$-th motor pool, and $\theta_i$ modulates the burst speed of the $i$-th synergy. The terms $\hat{\beta}_{ij}$ and $\hat{\theta}_i$ represent the corresponding baseline gains. The variable $p$ is the order of the term, with $n$ being the maximum order (set to four), and $\hat{\beta}_{ip}$ and $\hat{\theta}_{ip}$ are the coefficients associated with the $p$-th order for the $i$-th synergy. The parameter of $u$ is set to 1 for the gait speed of 1.3 $ms^{-1}$, the pace preferred by humans [6], [7]. The values of $u$ used for each speed are listed in Table S5. Note that $\hat{\beta}_{ip}$ is shared across all motor pools for a given synergy, i.e. it applies to all $\hat{\beta}_{ij}$ gains associated with the $i$-th synergy. Some specific $\theta$ gains are modulated by sensory afferents, reflecting known reflexive modulation of gait mechanics, following the formula:

$$\theta(t) = \hat{\theta}\frac{\lambda}{\lambda + F(t-\Delta t)} \tag{3}$$

where $\hat{\theta}$ is the baseline gain, $F$ is a specific sensory afferent, and $\lambda$ is a parameter that determines the modification of $\theta$ as a function of $F$. $F$ also undergoes sensory delay, $\Delta t$. Specifically, the $\theta$ of the *compliant-leg-behaviour synergy* (S1) is modulated by the positive force feedback from the vastus muscle ($F$ = VAS F+, $\Delta t_{10}$ = 10 ms), given the role of the knee extensors in compliant leg behaviour [2]. This modulation is common for the forward, backward and standing behaviours. For the *ankle push-off synergy* (S2), the positive force feedback from the soleus muscle ($F$ = SOL F+, $\Delta t_{20}$ = 20 ms) acts as sensory modulator, since this muscle controls plantarflexion at late stance [2]. This modulation only happens for forward and backward gait. The *transition synergy* (S6) is modulated by the forward leaning of the HAT ($F = \phi_{HAT}$, $\Delta t_5$ = 5 ms). $\phi_{HAT}$ is calculated with respect to the normal vector to the ground, with forward leaning taking positive values, and backward leaning taking negative values. Each synergy maintains the same $\lambda$ across all the internal models. The internal models also regulate the interconnections between synergies (Fig. S2). For forward gait, interconnections are set following a cyclical succession of synergies (S1→S2→S3→S4→S5). The last synergy, *leg-retraction* (S5) is followed by the *compliant-leg-behaviour* synergy (S1), closing the cycle. In backward gait, the positions of the *ankle push-off* synergy (S2) and the *leg-retraction* synergy (S5) are swapped (S1→S5→S3→S4→S2).

For forward and backward walking, two different connections are established between functional synergies according to their relative positions (Fig. S2). A strong inhibitory connection is established from one synergy to the synergy two positions behind (from $S_{i+2}$ to $S_i$):

$$S_i = \begin{cases} S_i & \text{if } S_{i+2} = 0 \\ 0 & \text{if } S_{i+2} \neq 0 \end{cases} \quad (4)$$

This mechanism avoids a functional synergy being still active when it is not needed anymore, and assures synchrony [8]. A second connection is established as a phase coupling [9]. Here, one functional synergy controls the rhythm of the previous synergy (from $S_{i+1}$ to $S_i$). The phase coupling is defined as:

$$\Phi_{mod} = -c \cdot \sin(\Phi_{i+1} + \Phi_i + \pi) \quad (5)$$

where $\Phi_{mod}$ is the phase interconnection between the synergies, $c$ is the strength of the coupling, manually tuned to 10, and $\Phi_i$ and $\Phi_{i+1}$ are the phases of the two consecutive functional synergies. The internal model of the standing task employs an auto-excitatory synapse of the *compliant-leg behaviour synergy* (S1) to achieve a tonic activation of the synergy. This dynamic is based on the phase of the same synergy:

$$S_1 = \begin{cases} \beta_{1j} & \text{if } \Phi_1 = 0 \\ \beta_{1j}\left(cos(\omega\theta\tau + \pi)\right) & \text{if } \Phi_1 \neq 0 \end{cases} \quad (6)$$

For each task, the same internal model is implemented separately in the right and left leg. This assures independent operativity, as the studies of the split-belt treadmill paradigm suggest [10].

#### E. *The Synergistic Layer*

The synergistic layer contains a vocabulary of six functional synergies (Table S2). Five synergies are dedicated to standing and gait and the sixth to transition between tasks. These numbers correspond with what is observed in literature, where five synergies are commonly identified for forward and backward gait [11], [12], [13], [14]. Each functional synergy is characterised by a discrete bell-shaped burst of activity (as suggested by Ivanenko et al. [12]) in the shape of a general equation:

$$A = \cos(\omega\tau + \pi) + 1 \quad (7)$$

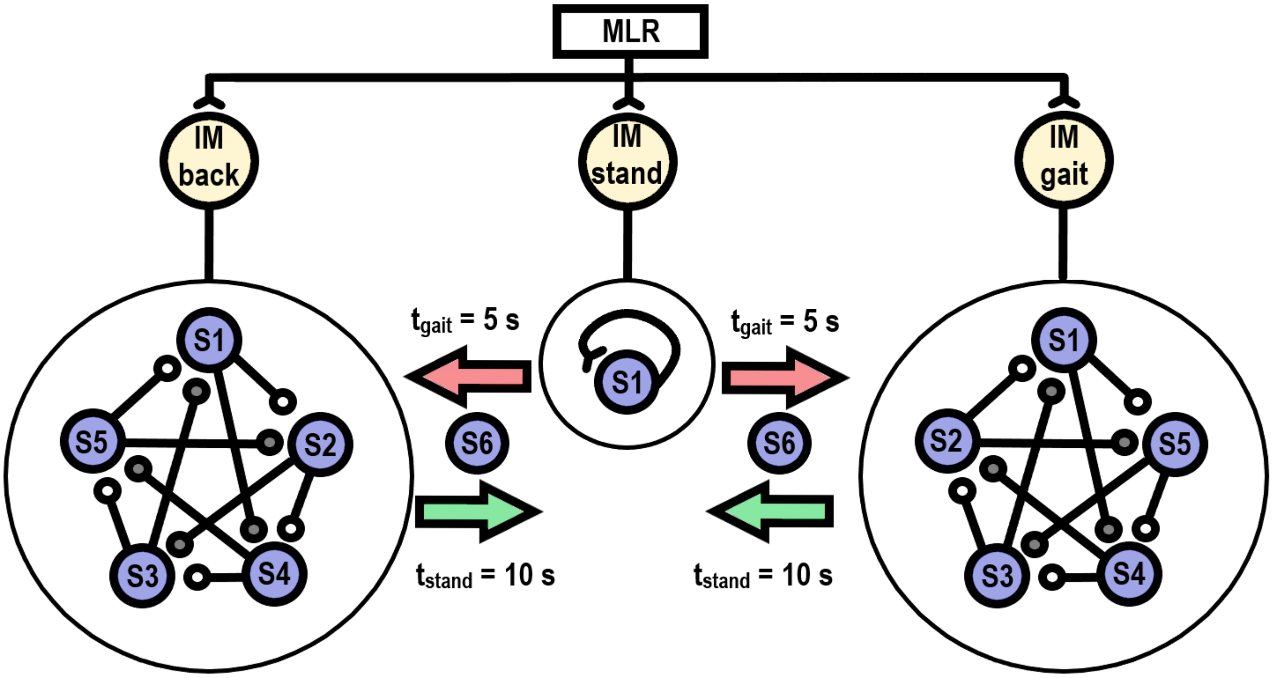


Figure S2. Flow of control of the IMMC from the MLR to the synergies. Three different networks are shown according to the controlling signal of the MLR, standing (middle), walking (right), and backward walking (left). Five different synergies are used to build these pathways. The sixth synergy accounts for all behaviour transitions, which are selected by the control signal. White interconnections withing the forward/backward internal models are the phase couplings, and the grey interconnections are the strong inhibitory synapses. The $\beta$ and $\theta$ gains and the lower layers are not indicated for simplicity. MLR: Mesencephalic Locomotor Region, IM: internal model, S1: compliant-leg behaviour, S2: ankle push-off, S3: hip unloading, S4: swing, S5: leg retraction, S6: transition.

where $\omega$ is angular frequency ($2\pi f$), and $\tau$ is the reference time for the synergy. The value of $f$ is set to 1 Hz by default. Synergies are triggered by different sensory afferents (Table S2) and their activation lasts until the angular frequency in Equation 7 ($\omega$) reaches $2\pi$. The parameter τ acts as an internal clock for the synergy, starting from zero every time the synergy is triggered. A trigger that occurs during a burst is ignored, so a burst always runs until its phase reaches $2\pi$. The exception is inhibition: when $S_{i+2}$ becomes active, the amplitude of $S_i$ is set to zero and its phase is advanced quickly to $2\pi$, so the burst ends with no further effect on the motor pools. The synergy general equation is modified by the internal models by adding θ gains and phase coupling, as in the case of forward and backward walking:

$$A_i = \cos(\omega\theta_i\tau + \pi + \Phi_i^{mod}) + 1 \tag{8}$$

$\Phi_{mod}$ is the value of phase interconnections between synergies, defined by the internal model according to Equation 5.
The general equation of the $i$-th synergy is modulated by the specific internal model, relating it to the $j$-th motor pool as:

$$S_{ij} = \beta_{ij}(cos(\omega\theta_i\tau + \pi + \Phi_{mod}) + 1) \tag{9}$$

where $\beta_{ij}$ is the spatial gain of the signal between the $i$-th synergy and the $j$-th motor pool. $\Phi_{mod}$ is the value of phase interconnections between synergies, defined by the internal model. Independently of the internal models, each functional synergy is defined by a function to accomplish and a specific trigger ($F_{thr}$):

**S1: Compliant-leg-behaviour.** This functional synergy resists gravity and avoids legs collapsing during standing or during heel strike in gait. This synergy is active during the stance phase [15] since it is triggered when the stance leg loads a weight higher than 20 N ($F_{thr}$ = *ipsilateral leg load* ≥ *20 N*).

**S2: Ankle push-off.** This functional synergy provides forward impulse and the acceleration of the trailing leg [16]. This functional synergy is triggered when the CoM reaches the leading leg ($F_{thr}$ = $CoM_x \geq heel_x$).

**S3: Hip unloading.** This functional synergy is active during late stance and assures that the leg unloads the weight of the body and prepares for swing. Hip flexion before swing correlates with extensor activity in the contralateral leg, implying that interlimb coordination must ensure the weight transfer to the leading leg [17]. Thus, the synergy is triggered when the load on the contralateral leg is higher than 20 N ($F_{thr}$ = *contralateral leg load* ≥ *20 N*).

**S4: Swing.** The purpose of this functional synergy is to provide the forward impulse of the swing leg to complete a step. This functional synergy is triggered when the ipsilateral leg is free of the body weight ($F_{thr}$ = *ipsilateral leg load* ≤ 20 N) [17] [18].

**S5: Leg retraction.** This functional synergy consists of a leg extension that opposes the swing motion and facilitates the foot landing. Bhounsule & Zamani [19] proposed that the swing-leg landing strategy depends on its kinematics during midstance. Considering midstance the instant when the gravity vector aligns with the leading leg, the synergy trigger is set to the heel of the swing leg passing over the CoM ($F_{thr}$ = $Heel_x \geq CoM_x$).

**S6: Transition.** This is a synergy whose role is to ease the transition by breaking quiet standing before forward and backward gait initiation, or to slow motion and place the feet together when transitioning back to stand. This functional synergy is not triggered by a sensorial event but by the activation of the corresponding control signals ($u > 0$).

The six functional synergies (Fig. S2) operate independently until an internal model organises them into a task-specific network. The same synergy can be employed in different networks, pursuing diverse motor objectives.

Table S2. List of functional synergies and their corresponding sensorial triggers and modulators.

| Functional synergy | | Trigger |
|---|---|---|
| 1) | Compliant leg behaviour | *Ipsilateral leg load* ≥ *20 N* |
| 2) | Ankle push-off | $COM_x \geq heel_x$ |
| 3) | Hip unloading | *Contralateral leg load* ≥ *20 N* |
| 4) | Swing | *Ipsilateral leg load* ≤ *20 N* |
| 5) | Leg retraction | $Heel_x \geq COM_x$ |
| 6) | Transition | *MLR control signal (u)* > 0 |

#### *F. The Motoneuron Layer*

The Motoneuron layer is constituted by the motor pools, which are agonist-antagonist groups of motoneurons that generate activation signals that are sent to the MTUs [20] [21]. The agonist-antagonist pairs are defined separately for each joint and each side of the body. One motor pool is dedicated to flexion and one to extension, so that the motoneuron layer of the IMMC is constituted by six motor pools per leg. Each monoarticular muscle receives input from a single motor pool. Monoarticular muscles are driven by the motor pool of the joint they cross: GLU by the hip extensor pool, HFL by the hip flexor pool, VAS by the knee extensor pool, BFSH by the knee flexor pool, SOL by the ankle extensor pool and TA by the ankle flexor pool. For biarticular muscles, two options were considered: scaling the activation of a muscle with a similar anatomical action, as in previous reflex-based models [1], [2], or summing the outputs of the two motor pools of the joints crossed. The choice was made heuristically. HAM and GAS receive a scaled copy of the activation of GLU and SOL, respectively, while RF receives the summed output of the hip flexor and knee extensor pools.

The activation output of the $j$-th motor pool ($y_j$) is defined as:

$$y_j = \sum_{i=1}^{n} S_{ij} + \sum_{k=1}^{m} \delta_k F_k(t - \Delta t) - \gamma y_{anta} \tag{10}$$

where $S_{ij}$ is the $i$-th functional synergy connecting to the $j$-th motor pool and $n$ is the number of synergies in the system. $F_k$ is the $k$-th reflex pathway stimulating that motor pool, $\delta_k$ is its related gain, m is the number of reflex pathways associated with the motor pool, and $\Delta t$ is the sensory delay affecting the sensory afferent (5, 10, or 20 ms). The inhibitory synapse from the antagonistic motor pool is denoted by $y_{anta}$, and $\gamma$ is its gain. As the antagonist motor pool also receives an inhibitory synapse, the pair is coupled by an agonist-antagonist disynaptic inhibition. Note that $S_i$ and $\delta_k$ can take positive or negative values, meaning that synergies and reflexes can inhibit each other (Fig. S3).

The reflexes maps to muscles acting on a specific joint through the corresponding motor pool. The contribution of reflex pathways to the motor pool is modulated by the corresponding $\delta$ gain. The hip motor pools are associated with two sensory afferents, mapping to their associated reflex pathways: the angular position, $F_{hip1} = \phi_{hip}$, and angular velocity of the hip, $F_{hip2} = \dot{\phi}_{hip}$, [22], each modulated by its own $\delta$ gain with a delay $\Delta t$ of 5 ms. For the knee, the reflex pathway inhibiting the extensor motor pool and exciting the flexor motor pool is knee pain, a nociception that measures overextension [1] [2]. Pain is equal to the knee angular position, $\phi_{knee}$, when it is lower than its neutral extended position, set to 0°. Also, this pathway is modulated by its own $\delta$ gain with a delay $\Delta t$ of 10 ms. The ankles present one reflex pathway which depends on two sensory afferents: the contact force at toes $f_{toes}$, and the contact force at heel, $f_{heel}$. These two sensory afferents are combined in a single reflex pathway which weights which side of the feet load more weight following the formula:

$$F_{ankle} = \frac{f_{toes}}{f_{toes} + f_{heel}} \tag{11}$$

This pathway is modulated by a single $\delta$ gain with a delay $\Delta t$ of 20 ms.

### G. *Optimisation and cost functions*

In this work we tested the IMMC through three simulation scenarios: Stand-To-Walk-To-Stand, Stand-To-Backward-Walk-To-Stand and forward walking with different speed transitions, starting from a standing position. In all the simulations the IMMC starts, at $t = 0$ s setting $u_{stand}$ to 1, thus activating the standing internal model. At an unspecified transition time, discovered during optimisation, the transition internal model is co-activated with the standing internal model, and at $t = 5$ s the transition and standing internal model are de-activated and the walking (forward or backward, depending on the task) internal model is activated. An equivalent procedure is applied when transitioning from walking to standing. The different internal models are characterised by different sets of $\beta$ and θ gains and by different connections between the synergies. The IMMC contains a total of 163 parameters to be optimized and 15 hand-tuned parameters, with some parameters shared across the different tasks and some that are task specific (see Table S3). The hand-tuned parameters are the θ gains of the standing and transition internal models (except for forward gait transition), the trigger thresholds, the synergy phase, the phase-coupling strength and the sensory delays. The optimisation is carried on in six stages (see Optimisation in the main text), each corresponding to a specific task, using the Covariance Matrix Adaptation Evolution Strategy [23].

The θ gains determine the relative temporal position of the synergies, and each synergy's burst frequency is calculated from its θ gain and that of the next synergy. For isolated synergies (e.g., in the standing and the transition internal models) two θ gains were used to compute the frequency despite only recruiting a single synergy. These were initially set to 0 and 0.25, respectively. In most cases, optimising them with the Covariance Matrix Adaptation Evolution Strategy was unnecessary, except for the forward gait transition task.

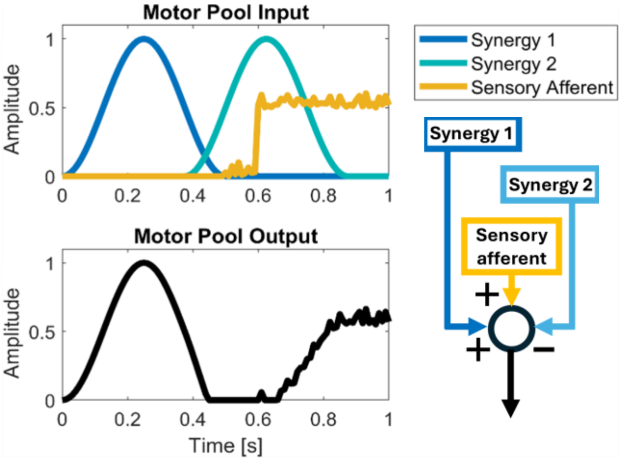


Figure S3. (Top) Hypothetical input to a motor pool, including two synergies and a sensory input. In this example, first synergy and sensory input have an excitatory synapse to the motor pool, while the second synergy has a negative synapse (Bottom) Resultant output of the motor pool. The sensory input is blocked by the second synergy.

Table S3. List of variables of the IMMC, which are optimised or hand-tuned, and which are shared between tasks.

| Variable | Symbol | Optimised | Hand-tuned | Shared |
|---|---|---|---|---|
| Spatial gains | $\beta, \hat{\beta}, \hat{\beta}_{ip}$ | x | | |
| Temporal gains | $\theta, \hat{\theta}, \hat{\theta}_{ip}$ | x | x | |
| Temporal gain modifiers | $\lambda$ | x | | x |
| Thresholds | $F_{thr}$ | | x | x |
| Sensory gain | $\delta$ | x | | x |
| Antagonist gain | $\gamma$ | x | | x |
| Synergy phases | $\Phi$ | | x | x |
| Phase coupling strength | $c$ | | x | x |
| Time delays | $\Delta t$ | | x | x |
| Transition times | $t_{trans}$, | x | | |

For each stage, the optimisation process used a dedicated cost function: standing $\left(J_{stand}\right)$, forward walking $\left(J_{gait}\right)$, forward gait speeds $\left(J_{speed}\right)$, backward walking $\left(J_{back}\right)$, and transitions from forward/backward walking back to standing $\left(J_{WTS-BWTS}\right)$. The cost function employed during stand optimisation, $J_{stand}$, tries to maximise time without falling while minimising energy consumption $(E)$:

$$J_{stand} = (t_{max} - t_{sim}) + E \tag{12}$$

where $t_{max}$ is the maximum time of simulation (5 s), and $t_{sim}$ is the time without falling. $E$ ($\mathrm{Wkg^{-1}}$) is the metabolic cost defined by Umberger et al. [24]. The cost function for forward gait ($J_{gait}$) is defined as:

$$J_{gait} = (t_{max} - t_{sim}) + 10\left|v_{tgt} - v_{sim}\right| + P + C \tag{13}$$

where $t_{max}$ is set to 10 s, $v_{tgt}$ is the objective speed of the simulation (for stage 2 equal to 1.3 $\mathrm{ms^{-1}}$), $v_{sim}$ is the average speed achieved during the simulation, $P$ is pain (the sum of joint torques when soft limits are exceeded) and $C$ ($\mathrm{Jkg^{-1}m^{-1}}$) is the metabolic cost of transport. Metabolic energy was included in the cost functions to favour economical solutions, as observed in human walking [6], [25]. It was computed for each muscle with the model of Umberger et al. [24] and summed over all muscles. $E$ is the resulting metabolic power normalised by body mass and averaged over the last 5 s of each simulated task (standing: t = 0–5 s; walking: t = 5–10 s; return to standing: t = 10–15 s). C is the metabolic energy over the same 5 s divided by body mass and by the distance travelled by the CoM. Walking speed was computed over the same window ($v_{sim}$ in Eqs. 13, 14, and 15). The parameters found in the 1.3 $\mathrm{ms^{-1}}$ walk optimisation were used as set values for the speed optimisations. In stage 3, only the coefficients of the speed modulation (Eqs. 1–2), which are common to all speeds, were optimised, with u set a priori for each speed (Table S5). All the forward speeds presented were included in the optimisation (stages 2 and 3).-The cost function ($J_{speed}$) is defined as:

$$J_{speed} = 35\left|v_{tgt} - v_{sim,last}\right| + C + \frac{AP}{3} \tag{14}$$

where $v_{tgt}$ is the target speed, $v_{sim,last}$ is the speed of the centre of mass in the anteroposterior axis in the last gait cycle to avoid excessive acceleration. The cost function for backward gait ($J_{back}$) is defined as:

$$J_{back} = (t_{max} - t_{sim}) + 10\left|v_{tgt} - v_{sim}\right| + \frac{AP}{2} + C \tag{15}$$

where $t_{max}$ = 10 s, $v_{tgt}$ is the target speed of the simulation (chosen to be 0.8 $\mathrm{ms^{-1}}$), and $v_{sim}$ is the average simulated speed. $AP$ is an ad-hoc cost function element that captures the deviation of the angular positions of the joints at heel strike with respect to their average. The cost function for the stand to walk transitions is identical for both transitions back to stand from forward and backward walking ($J_{WTS-BWTS}$) and is defined as:

$$J_{WTS-BWTS} = (t_{max} - t_{sim}) + 10\left|heel_{XR} - heel_{XL}\right| + E \tag{16}$$

where $t_{max}$ = 15 s is the total simulation time and $t_{sim}$ is the time completed without falling. The terms $heel_{XR}$ and $heel_{XL}$ represent the anteroposterior positions of the right and left heel and are used to evaluate the final distance between the feet in the recovered standing position.

#### *H. Rationale for the architecture*

The organisation of the IMMC follows from three observations in the modularity literature. Standing, forward walking and backward walking can be reconstructed from largely the same muscle synergies [11], [14], [26], which motivated a single vocabulary of functional synergies shared by all tasks. Locomotor synergies are expressed as brief, phase-specific bursts [12], [13], and stepping depends on load and limb-position afferents [17], [18], so each synergy was implemented as a bell-shaped burst triggered by a sensory event. Differences between tasks appear mainly in the timing and weighting of the shared synergies [14], [26], and these were assigned to task-specific internal models (β, θ and synergy ordering). Assigning each locomotor synergy to one principle of legged mechanics [2] gave it a defined mechanical function and a natural trigger. Table S4 lists, for each element of the architecture, its experimental basis and the modelling choices made in its implementation.

Table S4. Experimental basis and modelling choices for each element of the IMMC.

| **Element** | **Experimental basis** | **Modelling choice** |
|---|---|---|
| Shared synergy vocabulary | Same synergies, with different timing, in forward and backward walking [11], [14]; common synergies for balance and walking [26], [27] | Six synergies: five shared by standing and gait, one added for transitions; backward walking swaps S2 and S5 in the cyclic sequence |
| Function of S1–S5 | Principles of legged mechanics emerge in reflex-driven walking [2] | One-to-one mapping of each synergy to one principle |
| Burst waveform | Impulsive, phase-specific locomotor activation [12], [13] | Raised-cosine burst, 1 Hz default (Eq. 7) |
| Sensory triggers | Load and hip-position afferents gate stepping and swing onset [17], [18]; midstance kinematics shape landing [19] | Choice of trigger signals and thresholds (Table S2) |

| | | |
|---|---|---|
| Sensory modulation of burst duration | Positive force feedback during stance [2], [15] | θ modulated by VAS and SOL force and HAT lean (Eq. 3) |
| Transition synergy (S6) | Introduced in this work | Ad hoc synergy activated by MLR commands |
| Synergy interconnections | Introduced in this work | Inhibition from S_{i+2} and phase coupling (Eqs. 4–5), adapted from network models [8], [9] |
| Internal models | Task-specific spinal interneuron pathways [28], [29] | IM as a task-specific set of β, θ and connectivity acting on shared synergies |
| Independent left/right IMs | Independent control of each leg in split-belt adaptation [10] | One copy of each IM per leg |
| Control layer (MLR) | MLR initiates locomotion and scales its speed [5], [30] | Discrete signals select IMs; speed set through polynomials of u (Eqs. 1–2) |
| Motor pools | Spinal motor pools [20], [21]; reciprocal inhibition between antagonists | One flexor and one extensor pool per joint with mutual inhibition γ (Eq. 10) |
| Reflex pathways | Hip, knee and ankle feedback used in reflex gait models [1], [2], [22] | Pathways and gains as in Eqs. 10–11 |
| Sensory delays | Conduction-distance delays used in previous models [2], [3] | 5, 10 and 20 ms by joint |

#### I. *Data analysis*

Simulated data were segmented into gait cycles between consecutive heel strikes of the same leg and normalised to 100 % of the gait cycle. To exclude gait initiation, the first 1.5 s of forward walking (until $t = 6.5$ s) and the first 1 s of backward walking (until $t = 6$ s) were discarded. In the speed simulations, the control signal changes to its target value at $t = 6.5$ s, and the same initial period was discarded. The analysis used ten gait cycles (five per leg) for forward walking at 1.3 ms$^{-1}$ and for backward walking, and ten cycles on average for the other forward speeds. For the experimental data, ten gait cycles were selected starting from the middle of each recording and segmented in the same way.

Muscle activations were normalised before the comparison. The minimum of each experimental activation was subtracted, and the result was rescaled to the maximum of the original activation. The minimum of each simulated activation was also subtracted, and the result was scaled so that its maximum matched that of the corresponding experimental activation. Simulated and experimental activations therefore have the same peak, so the comparison in Table 1 of the main text reflects differences in shape and timing, and peak magnitude differences are not reported for the muscles. The cross-correlation coefficient ($r$) and its lag (Δ) were computed between the averaged simulated and experimental traces, together with the root-mean-square error (RMSE), the difference between their maxima (peak magnitude difference) and the difference in the gait-cycle percentage at which the maxima occur (peak timing difference).

### II. Supplementary Results

#### A. *Speed transitions biomechanics*

In the speed transitions simulations, the angular positions of the joints in the lower (0.8 ms$^{-1}$) and upper (1.8 ms$^{-1}$) walking speed simulations were compared to the experimental data of subjects walking at slow and fast speeds, respectively derived from a publicly available dataset [31] (Fig. S4(A)). The cross-correlation coefficients ($r$) for hips and knees are 0.90 ($\Delta$ = 0 %) and 0.98 ($\Delta$ = 0 %) at slow speed, and 0.86 ($\Delta$ = 0 %) and 0.97 ($\Delta$ = 1 %) at fast speed. The ankle shows a cross-correlation below 0.7 at 0.8 ms$^{-1}$ ($r_{ankle,0.8}$ = 0.56, $\Delta$ = -5 %), but above 0.7 at 1.8 ms$^{-1}$ ($r_{ankle,1.8}$ = 0.74, $\Delta$ = -4 %). The ankle behaviour for both speeds is generally comparable with that observed for the 1.3 ms$^{-1}$ simulations (Figure 4(A)). The joint trajectories of the other simulations (speeds of 0.9, 1.2, 1.5, and 1.6 ms$^{-1}$) were also compared to the corresponding slow or fast speeds exhibited by subjects (Table S5). Except for the ankle angular positions at 0.9 and 1.5 ms$^{-1}$, all the trajectories presented a good match ($r > 0.7$). The GRFs (Fig. S4(B)) for the 0.8 ms$^{-1}$ and 1.8 ms$^{-1}$ speed simulations are similar to the experimental samples [31], which is supported by the cross correlation analysis ($r_{GRF,0.8}$ = 0.95, $\Delta$ = -2 %; $r_{GRF,1.8}$ = 0.85, $\Delta$ = -2 %). The cross-correlation coefficients for the rest of speed transitions are above 0.7 (Table S5). The cost of transport (Fig. 3 of the main text) was lowest at 1.3 and 1.5 ms$^{-1}$ ($u$ = 1 and 0.8; Table S5), close to the value at which the forward-walking internal model was optimised ($u$ = 1), and increased as $u$ moved away from these values. Fig. S4(C) shows the simulated muscle activation timing compared with experimentally observed normative [32]. Also in this case, simulated patterns of activation are comparable to experimental ones, except for the RF, GLU and HAM for the 1.8 ms$^{-1}$. Foot-off event is compared between experimental one [14] (67 %) with the simulated foot-off at 0.8 ms$^{-1}$ (67 %) and 1.8 ms$^{-1}$ (71 %). Video STW-WTS.mp4 shows the complete 0.8 ms$^{-1}$ simulation and video forward_speeds.mp4 (1.8 ms$^{-1}$) shows the complete 1.8 ms$^{-1}$ simulation.

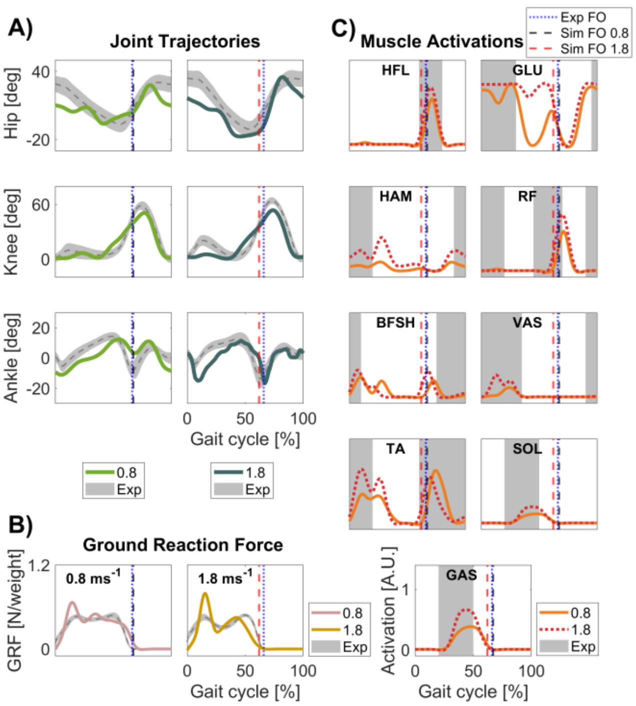


Figure S4. (A) Experimental and simulated angular positions for the lower (0.8 ms-1) and the upper (1.8 ms-1) transitions. Light green: simulated data at 0.8 ms-1, dark green: simulated data at 1.8 ms-1, grey: experimental data. Upward direction: flexion, downward direction: extension. (B) GRF for left foot along the duration of walking for the lower (0.8 ms-1) and the upper (1.8 ms-1) transition. Pink: simulated data at 0.8 ms-1, yellow: simulated data at 1.8 ms-1, grey: experimental data. (C) Muscle activation patterns for the left and right leg along the gait cycle for the lower and the upper transitions. Light orange solid line: simulated data at 0.8 ms-1, dark orange dotted line: simulated data at 1.8 ms-1, grey: experimental data. Experimental angular position and GRF are given by Fukuchi et al. [31]. Muscle activation timing is interpreted from Zajac et al. [32] .Blue vertical dotted line: generic experimental foot-off (67 %), black vertical dotted line: simulation foot-off at 0.8 ms-1 (67 %), red vertical dotted line: simulation foot-off at 1.8 ms-1 (71 %).

Table S5. Control-signal value ($u$) for each forward walking speed. Cross-correlation coefficients and lags of the simulation and the experimental [31] joint angular positions. The 1.3 ms$^{-1}$ simulation is compared with the data of [14] and is detailed in Table 1 of the main text.

| Speed [ms$^{-1}$] | $u$ | $r$ (hip/knee/ankle/GRF) | lag [%] (hip/knee/ankle/GRF) |
|---|---|---|---|
| 0.8 | 1.6 | 0.90/0.98/0.56/0.95 | 0/-2/-5/-1 |
| 0.9 | 1.4 | 0.90/0.98/0.59/0.93 | 0/-4/-5/0 |
| 1.2 | 1.2 | 0.88/0.99/0.78/0.94 | 0/-2/-4/1 |
| 1.3 | 1 | 0.92/0.96/0.84/0.99 | 0/-3/-2/0 |
| 1.5 | 0.8 | 0.89/0.97/0.69/0.91 | 0/-2/-5/-1 |
| 1.6 | 0.6 | 0.88/0.97/0.76/0.91 | 0/1/-5/2 |
| 1.8 | 0.4 | 0.86/0.97/0.74/0.85 | 0/1/-4/1 |

#### *B. Transitions, speed settling and longer simulations*

Fig. 2 of the main text shows the onset of the transition internal model, the switches to gait and to standing, the first heel strike after gait initiation and the last foot contact before standing (A, B), and the stride-averaged speed of each forward simulation (C), computed between consecutive heel strikes of the same foot. All speed simulations share the same gait initiation and diverge when $u$ is set to its target value at $t$ = 6.5 s. Table S6 reports, for each speed, the speed of the last stride, the mean stride speed and the settling time, defined as the first time after which the stride-averaged speed stays within ±5 % of its final value. In the original 10-s simulations, the mean stride speed was within 8 % of the target speed, but at 1.2 ms$^{-1}$ and above the stride speed was still increasing at the end of the simulation, and the settling times (8.7–9.7 s) were close to its end.

Table S6. Speed of the last stride, mean stride speed, settling time (first time after which the stride-averaged speed stays within ±5 % of its final value) and end time of the original and extended simulations. B: backward walking.

| Target speed [ms$^{-1}$] | 0.8 | 0.9 | 1.2 | 1.3 | 1.5 | 1.6 | 1.8 | 0.8 B |
|---|---|---|---|---|---|---|---|---|
| *Original simulations (10 s)* | | | | | | | | |
| Last-stride speed [ms$^{-1}$] | 0.86 | 0.95 | 1.52 | 1.67 | 1.80 | 2.08 | 2.24 | 0.99 |
| Mean stride speed [ms$^{-1}$] | 0.83 | 0.90 | 1.16 | 1.26 | 1.38 | 1.53 | 1.69 | 0.86 |
| Settling time [s] | 6.52 | 7.30 | 9.66 | 9.60 | 8.84 | 8.87 | 8.72 | 5.76 |
| End of simulation [s] | 10.00 | 10.00 | 10.00 | 10.00 | 10.00 | 10.00 | 10.00 | 10.00 |
| *Extended simulations (up to 30 s)* | | | | | | | | |
| Last-stride speed [ms$^{-1}$] | 0.76 | 1.63 | 1.71 | 1.74 | 1.84 | 2.04 | 2.11 | 0.93 |
| Mean stride speed [ms$^{-1}$] | 0.76 | 1.34 | 1.48 | 1.54 | 1.68 | 1.90 | 1.87 | 0.79 |
| Settling time [s] | 6.60 | 13.63 | 12.70 | 12.62 | 10.44 | 9.73 | 8.05 | 8.18 |
| End of simulation [s] | 11.19 | 19.50 | 19.69 | 19.56 | 19.78 | 19.88 | 19.94 | 11.19 |

To assess the behaviour over longer periods, the simulations were extended up to 30 s (Fig. S5). The longest simulations completed without falling ended at about 20 s for the forward speeds between 0.9 and 1.8 ms$^{-1}$, and at about 11 s at 0.8 ms$^{-1}$ and in backward walking. Between 0.9 and 1.8 ms$^{-1}$, the forward speed increased until 8–14 s and then settled at 1.6–2.1 ms$^{-1}$, above the target speeds, whereas forward walking at 0.8 ms$^{-1}$ and backward walking settled at 0.76 and 0.93 ms$^{-1}$, respectively. These durations exceed the 10 s used to optimise gait, and the controller has no feedback on walking speed, which may explain the drift.

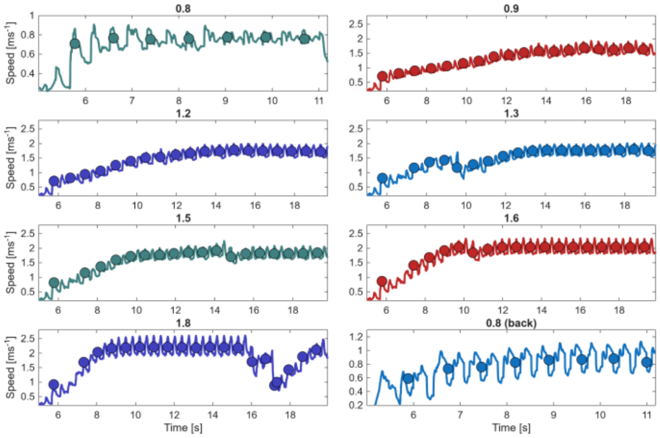


Figure S5. Instantaneous HAT speed (lines) and stride-averaged speed (circles) in the extended simulations of forward walking at each speed and of backward walking at 0.8 ms$^{-1}$, each shown up to the longest duration completed without falling.

Table S7 reports the outcome of the transition simulations when $t_{gait}$ and $t_{stand}$, together with the corresponding onsets of the transition internal model, were shifted by ±0.25 s and ±0.5 s. The forward transitions were completed in all simulations in which gait was commanded earlier than in the optimised schedule, regardless of the shift of $t_{stand}$, and failed in all those in which it was commanded later. The backward transitions were completed only when both commands were advanced by 0.5 s. The standing internal model was optimised over 5 s, which may explain why delaying the gait command, and thus prolonging quiet standing, led to failure.

Table S7. Outcome of the Stand-To-Walk-To-Stand (forward) and Stand-To-Backward Walk-To-Stand (backward) simulations when $t_{gait}$ and $t_{stand}$ were shifted with respect to the optimised schedule; the onsets of the transition internal model were shifted by the same amount. Yes: transition to gait and back to standing completed without falling.

| Shift of $t_{gait}$ / $t_{stand}$ [s] | Forward | Backward |
|---|---|---|
| +0.25 / +0.25 | No | No |
| −0.25 / −0.25 | Yes | No |
| +0.5 / +0.5 | No | No |
| −0.5 / −0.5 | Yes | Yes |
| +0.25 / −0.25 | No | No |
| −0.25 / +0.25 | Yes | No |
| +0.5 / −0.5 | No | No |
| −0.5 / +0.5 | Yes | No |

### C. *Muscle synergies*

Muscle synergies were extracted by non-negative matrix factorisation (NMF) from the simulated and experimental activations of forward walking at 1.3 $ms^{-1}$ and backward walking at 0.8 $ms^{-1}$, using the average gait cycles analysed in Table 1 of the main text (Section I). The analysis included the eight muscles recorded experimentally (HFL, GLU, RF, BF, VAS, TA, SOL and GAS), with the simulated BFSH compared with BF; HAM was excluded. For each data set, NMF was repeated 50 times from random initial values and the solution with the lowest reconstruction error was kept. Three synergies were extracted, the smallest number that accounted for at least 90 % of the variance (VAF) in all four data sets (Table S8). Each simulated synergy was paired with an experimental one so as to maximise the total cosine similarity of the weights, and the activation coefficients of each pair were compared with Pearson's correlation ($r$).

In forward walking, each experimental synergy had a simulated counterpart (Fig. S6, Table S8): loading (GLU, VAS and BF; cosine similarity 0.90, $r$ = 0.72), ankle push-off (SOL and GAS; 0.94, $r$ = 0.94) and swing (HFL, with TA and RF; 0.93, $r$ = 0.54). The simulated loading synergy had a second peak at about 30 % of the gait cycle, and the simulated swing burst was shorter than the experimental one. In backward walking, the simulated synergies matched the experimental synergy of VAS, TA and RF (0.96, $r$ = 0.96). The other two experimental synergies, which combine SOL or GAS with hip muscles, were matched only in part (0.61 and 0.65): the simulated synergies contained the hip muscles but not SOL and GAS, which are not excited in the simulated backward gait (Fig. 5 of the main text).

Table S8. Muscle synergies of forward walking at 1.3 $ms^{-1}$ and backward walking at 0.8 $ms^{-1}$. For each experimental synergy (numbered as in Fig. S6), the table gives its main muscles, the cosine similarity between its weights and those of the paired simulated synergy, and the correlation ($r$) between their activation coefficients. VAF: variance accounted for by three synergies.

| Synergy | Main muscles (experimental) | Cosine similarity | $r$ |
|---|---|---|---|
| *Forward walking, 1.3 $ms^{-1}$ (VAF: experimental 0.95, simulated 0.93)* | | | |
| 1 | GLU, VAS, BF | 0.90 | 0.72 |
| 2 | SOL, GAS | 0.94 | 0.94 |
| 3 | HFL, TA, RF | 0.93 | 0.54 |
| *Backward walking, 0.8 $ms^{-1}$ (VAF: experimental 0.93, simulated 0.96)* | | | |
| 1 | VAS, TA, RF | 0.96 | 0.96 |
| 2 | SOL, GLU, HFL | 0.61 | 0.76 |
| 3 | GAS, BF, HFL | 0.65 | 0.77 |

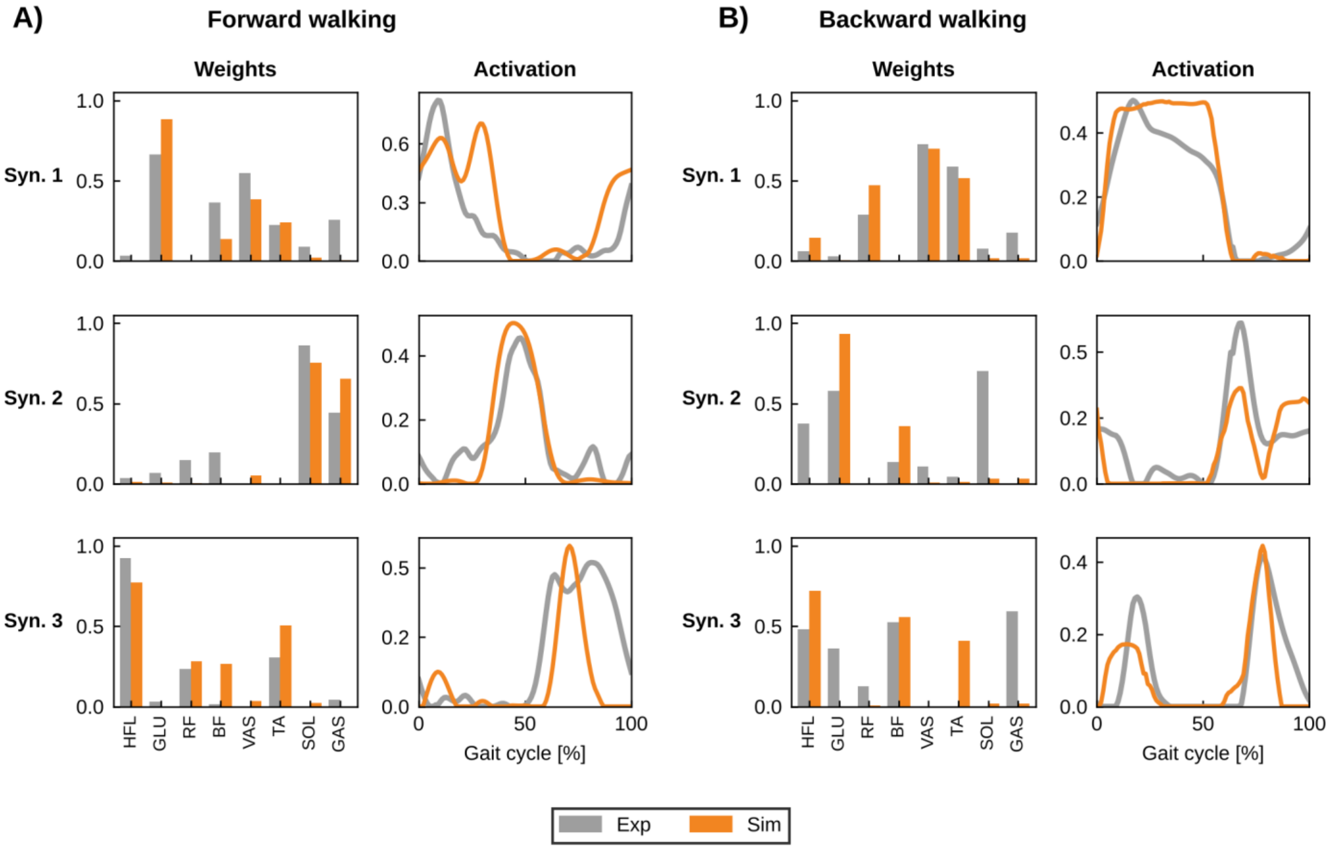


Figure S6. Muscle synergies extracted from the experimental (grey) and simulated (orange) muscle activations in (A) forward walking at 1.3 ms⁻¹ and (B) backward walking at 0.8 ms⁻¹. For each synergy, the left panel shows the muscle weights and the right panel the activation coefficient over the gait cycle. Each simulated synergy is shown with the experimental synergy it was paired with (Table S8). For the simulation, BF denotes BFSH.
